# Influence of short and long term processes on SAR11 communities in open ocean and coastal systems

**DOI:** 10.1101/2022.06.17.496560

**Authors:** Luis M. Bolaños, Karen Tait, Paul J. Somerfield, Rachel J. Parsons, Stephen J. Giovannoni, Timothy Smyth, Ben Temperton

## Abstract

SAR11 bacteria dominate the surface ocean and are major players in converting fixed carbon back to atmospheric carbon dioxide. The SAR11 clade is comprised of niche-specialized ecotypes that display distinctive spatiotemporal transitions. We analysed SAR11 ecotype seasonality in two long-term 16S rRNA amplicon time series representing different North Atlantic regimes: the Sargasso Sea (subtropical ocean-gyre; BATS) and the temperate coastal Western English Channel (WEC). Using phylogenetically resolved amplicon sequence variants (ASVs), we evaluated seasonal environmental constraints on SAR11 ecotype periodicity. Despite large differences in temperature and nutrient availability between the two sites, at both SAR11 succession was defined by summer and winter clusters of ASVs. Summer cluster was dominated by ecotype Ia.3 in both sites. Winter clusters were dominated by ecotypes Ib and IIa.A at BATS and Ia.1 and IIa.B at WEC. A two-year weekly analysis within the WEC time series showed that the response of SAR11 communities to short-term environmental fluctuations was variable. In 2016, community shifts were abrupt and synchronised to environmental shifts. However, in 2015, changes were gradual and decoupled from environmental fluctuations, likely due to increased mixing from strong winds. We demonstrate that interannual weather variability disturb the pace of SAR11 seasonal progression.

## Introduction

SAR11 are heterotrophic, free-living planktonic alpha-proteobacteria with streamlined genomes and small cell size (~0.04 μm^3^) (1–4). SAR11 are globally distributed throughout the surface oceans and account for around 25% of the plankton cells in this layer of the water column (5–7) At a broad phylogenetic level, SAR11 order (Pelagibacterales (8)) is divided in subclades that are congruent with spatial and temporal distributions defined by environmental factors such as depth, latitude, and season (9–12). Therefore, these subclades have been defined as ecotypes (11). Genomic studies have determined clear phylogenetic subclade boundaries within SAR11 clade and around half of these lack a cultivable representative (13). While fundamentally similar in phenotype and core genome conservation, SAR11 ecotypes display differences in many characteristics that have adaptive and geochemical significance. Variation among SAR11 ecotypes in phosphorous compound metabolism (14), carbon utilization (15,16), production of gaseous compounds (17) and utilization rates of volatile organic compounds (18) have important implications for ocean (19,20) and atmosphere (21) geochemistry.

16S rRNA surveys across oceanic environmental gradients have shown that SAR11 communities differ by favouring the most adapted ecotypes (22,23). Time series in temperate regions of the ocean revealed that SAR11 communities are sensitive to seasonal changes, which determine the successional progression of dominant ecotypes (11,24–28). Specifically, long-term time series have shown that SAR11 dynamics display a highly consistent ecotype periodicity through seasons (27,29). Short time-scale variation (days to weeks) of SAR11 using high frequency sampling suggests that SAR11 communities gradually transition through the seasons (30), yet the influence of short-term environmental fluctuations on SAR11 variability remains unexplored.

In the North Atlantic western subtropical gyre, the Bermuda Atlantic Time-series Study (BATS) have collected oceanographic and biological data since 1988 (31). 16S rRNA surveys have been consistently generated within these years from monthly sampling (11,29). The Western English Channel (WEC), Station L4, located off the southwestern coast of the United Kingdom is one of the longest oceanographic time series of the world (32). Since the year 2000, samples from the WEC have been collected weekly for multiple microbial molecular studies (33–36). These time series provide a robust framework to quantify the influence of environmental factors on SAR11 composition across long and short time scales, and identify similarities and differences between temperate coastal waters and the subtropical open ocean.

We analysed the SAR11 fraction of the 16S rRNA surveys retrieved from datasets spanning more than 27 years since 1991 in BATS and seven continuous years (2012-2018) in the WEC. SAR11 amplicon sequence variants (ASV) were phylogenetically classified using PhyloAssigner pipeline (29) and a curated phylogenetic database of full-length SAR11 16S rRNA gene sequences (22). This approach allows robust comparisons between different sequencing technologies because it relies on maximum likelihood placement of reads on a pre-determined phylogeny. In both time series, we evaluated the persistence of SAR11 ecotype succession and the influences of the environment on the annual progression of SAR11 composition. Finally, we leveraged the near-complete, weekly sampling within a two-year period in the WEC to evaluate the short-term environmental influences within the annual progression. The results provide a panoramic overview of SAR11 responses to interannual fluctuations in the surface ocean.

## Materials and Methods

### Sample collection, DNA extraction, 16S rRNA library preparation, and sequencing

#### Bermuda Atlantic Time-series

Depth profiles were collected monthly at BATS (31°40′N,64°10′W). Surface samples (1-5m) for the periods 1991-2002 (29) and 2016-2018 were analysed in this study. Microbial biomass from four litres of water was collected on 47 mm, 0.22 μm pore Supor filters (Sigma-Aldrich, St. Louis, MO, USA) from 1991 to 2002 and on Sterivex filters of the same composition from 2016 to 2018. DNA was extracted using a phenol:chloroform protocol (37–39). 454 FLX sequences were retrieved from iMicrobe (CAM_P_0000950). 16S rRNA V1-V2 amplicons from samples collected between 2016 and 2018 were generated with the primers 27F (5′-AGAGTTTGATCNTGGCTCAG-3) (40) and 338RPL (5′-GCWGCCWCCCGTAGGWGT-3′) (29,41). Amplicon libraries (2016-2018) were prepared with the Nextera XT Kit (Illumina Inc.). Libraries were sequenced using the MiSeq platform (v.2; 2×250) by the Centre for Genome Research and Biocomputing (Oregon State University).

#### Western English Channel

Surface water samples (1-5m) were collected weekly at the station L4 (50°15′N,4°13’W) as part of the Western Channel Observatory time series. Five litres of water were filtered through 0.22 μm pore Sterivex filters. DNA was extracted using a Qiagen AllPrep DNA/RNA Micro Kit. 16S rRNA V4-V5 amplicons were generated with the primers 515fB (5′-GTGYCAGCMGCCGCGGTAA-3′) and 806rB (5’-GGACTACNVGGGTWTCTAAT-3’). Amplicon libraries were generated with the Nextera XT Kit (Illumina Inc.). Libraries were sequenced using the MiSeq platform (v.3; 2×300) by NU-OMICS (Northumbria University, U.K.). A list of samples is provided as Table S1.

### Sequence processing and taxonomic classification

Surface 454 FLX amplicon datasets from BATS spanning from 1991 to 2002 (29) and the MiSeq Illumina datasets (2015-2018 BATS and 2012-2018 WEC) were quality filtered, dereplicated and merged with Dada2 (42) and phyloseq (43) R packages. SAR11 sequences were extracted, aligned and phylogenetically placed on a 16S rRNA full-length custom tree (22) with Phyloassigner v6.166 (29) (see Supplementary methods).

### SAR11 composition and seasonality analysis

Relative contribution, rarefaction, β diversity, ordinations, and Euclidean distances were calculated in R v4.0 (44) using phyloseq v1.34 (43), vegan v2.5 (45) and ggplot2 v3.3 (46) (see Supplementary methods). Figures were edited in Inkscape2 (www.inkscape.org) for aesthetics. Seasonality, defined as periodic changes of ASVs, was determined with a Fisher G-test (47) implemented in GeneCycle R package (48) based on relative contributions.

### Temporal diversity and influence of environmental variables on SAR11 composition

β diversity estimated by the Bray–Curtis dissimilarity index between all samples were used to examine the annual cycles of SAR11 communities in both sites throughout the long-term sampling. To evaluate the influence of the environmental variability on SAR11 composition within short-time scales, we performed a wavelet coherence analysis with R package WaveletComp (49) on the weekly WEC sampling from April 2015 to April 2017 (see Supplementary methods).

## Results

### SAR11 ecotype seasonality is persistent through multiannual time series in the Sargasso Sea and Western English Channel

At BATS, SAR11 comprised between 4.4% to 65.4% of the community (Fig. 1a). Sequenced with 454 FLX, SAR11 relative contribution from 1991 to 2004 at BATS showed a broad distribution of values (29.7±12.8%). However, the period from 2016 to 2018 (Illumina;MiSeq) displayed a constant high contribution with more than 85% samples >40% SAR11, with a narrower distribution (45.9±8.5%). This difference could be explained by the sequencing saturation achieved by the higher coverage of Illumina datasets compared to those generated by 454 FLX. SAR11 relative contribution was more variable within the WEC time series (Illumina;MiSeq), ranging from 0.04% to 53.9% (15.7±9.7%) (Fig. 2a) with a decrease in SAR11 contribution during summer months (June, July, August). This result supports previous observations of SAR11 seasonality at WEC (34,50).

**Figure 1:**
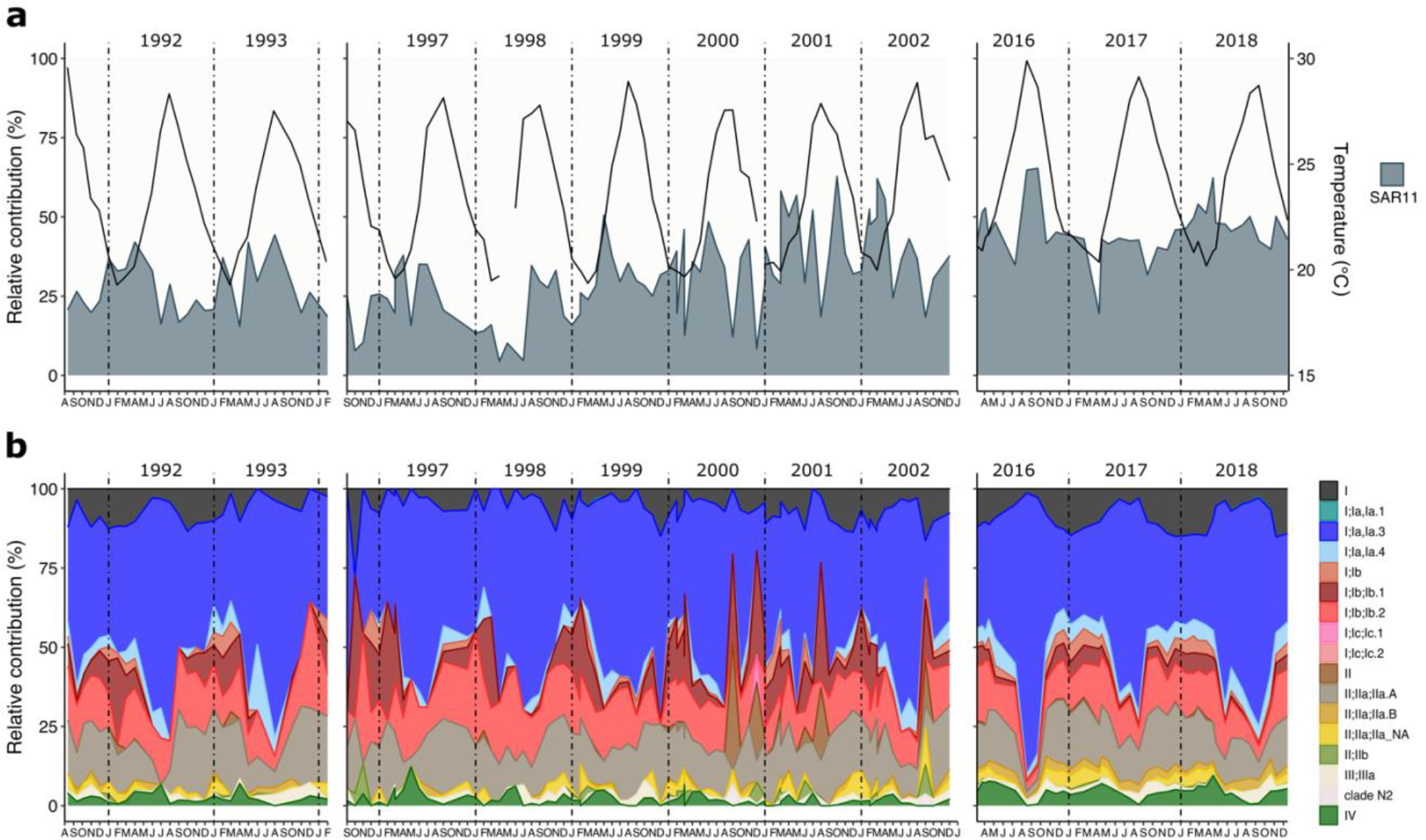
Monthly relative contribution of surface SAR11 ecotypes through multiannual 16S rRNA surveys at the Bermuda Atlantic Time Series. **(a)** Monthly relative contribution of the SAR11 fraction to the total of the amplicon dataset. The black line represents temperature through the sampled years. **(b)** Monthly relative contribution of ecotypes to the total SAR11 fraction. Samples from 1991-1994 and 1996-2002 were sequenced using 454 FLX technology, while 2016-2018 were sequenced using the Illumina MiSeq platform.

**Figure 2:**
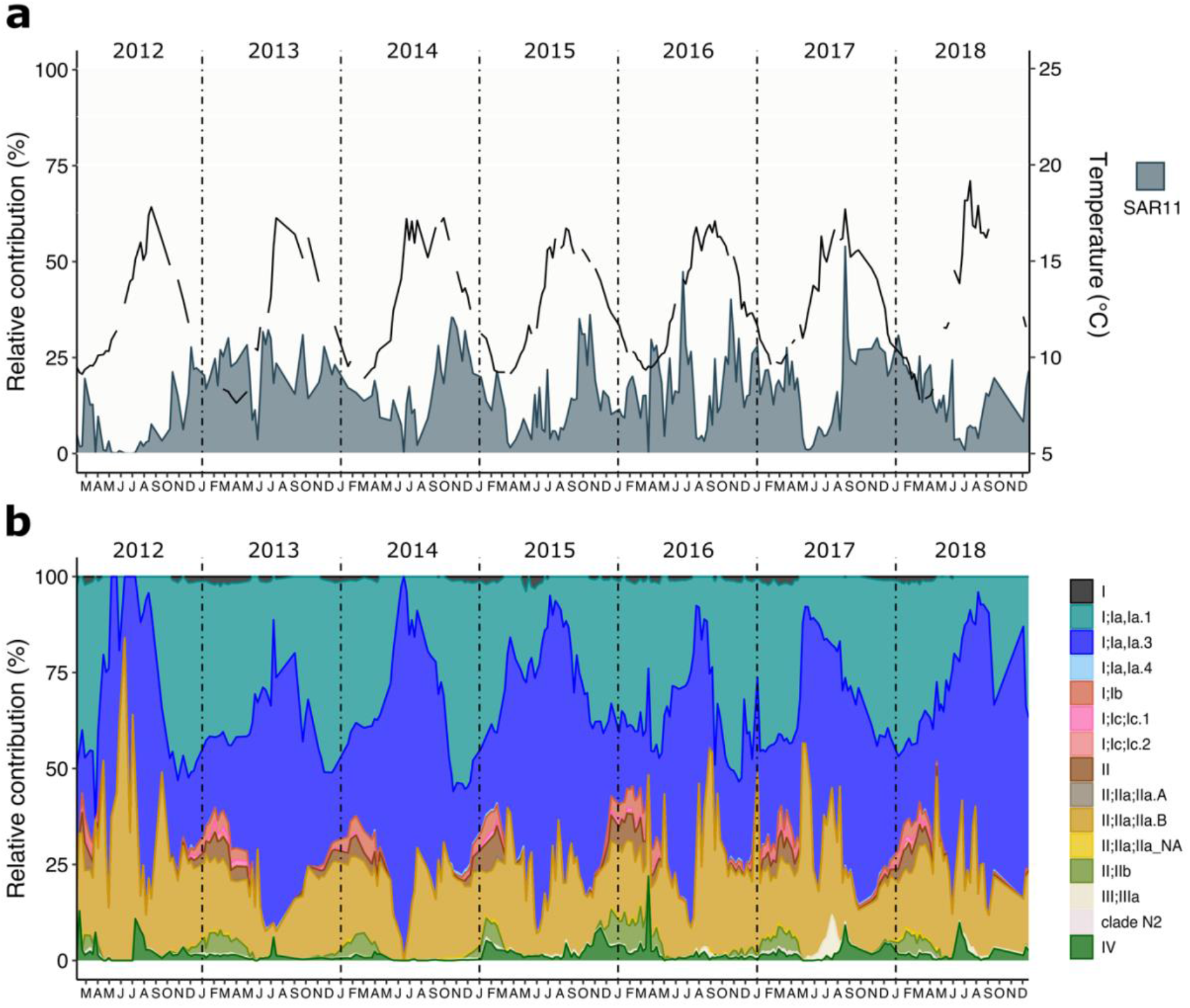
Monthly relative contribution of surface SAR11 ecotypes through a multiannual 16S rRNA survey at the station L4 from the Western English Channel. **(a)** Monthly relative contribution of the SAR11 fraction to the total of the amplicon dataset. The black line represents temperature through the sampled years. **(b)** Monthly relative contribution of ecotypes to the total SAR11 fraction.

SAR11 Ia ecotypes dominate the surface of both locations, comprising 49% of the total at BATS (Fig. 1b,S1) and 71.9% at WEC (Fig. 2b,S2). Ecotype Ia.3 (“warm water” (12)) dominates at BATS, showing an oscillatory progression reaching a maximum during the stratified summer and decreasing to the lowest contributions in the deepest mixing of the water column in winter, thereafter, returning to an increasing progression. Unexpectedly, given the comparatively low seasonal temperatures [BATS maximum 29.6 °C, minimum 19.3 °C; WEC maximum 19.2 °C, minimum 7.7 °C], ecotype Ia.3 ASVs were also highly abundant in the WEC, comprising 35.8% of SAR11 sequences, and displayed a similar annual pattern to that observed in BATS. Ecotype Ia.1, prominent in cold regions (12), was negligible at BATS as expected (Figs. 2b,S3). At WEC, Ia.1 made up 35.9% of SAR11 sequences and displayed an opposite oscillation pattern to Ia.3, with an increasing progression starting in summer (June-July), reaching a peak in early winter (December-January). Ecotype IIa.B was the third most abundant in WEC (18.8%). It displayed a similar annual progression to Ia.1, reaching a maximum peak in winter. Other ecotypes that increased its relative contribution during winter were Ib, II, and IIb. Clade IV displayed low contributions with periodic peaks after summer. At BATS, the progression to winter was dominated by Ib ecotypes and IIa.A (instead of IIa.B at WEC). Additionally, Ia.4, IV, non-determined I and IIa accompanied it. These differences of winter dominant ecotypes between the two time series might be related to adaptations to the different temperature range and nutrient concentrations intrinsic of each environment (Figs. S4,S5,S6).

SAR11 communities showed a recurrent annual cycle in both locations (Fig. 3). Pairwise comparison of Bray-Curtis dissimilarities of rarefied SAR11 ASVs over time did not show a clear annual pattern from 1991 to 2002 in the 454 FLX data (Fig. 3a), most likely as a result of reduced coverage of the community. In contrast, the 2016-2018 datasets (Illumina) showed a consistent sinusoidal pattern with a wavelength of one year (Fig. 3b). In the WEC, a similar one-year sinusoidal pattern was also evident, with greater clarity provided by the weekly sampling. Community turnover was maximised approximately every 180 days in the WEC data.

**Figure 3:**
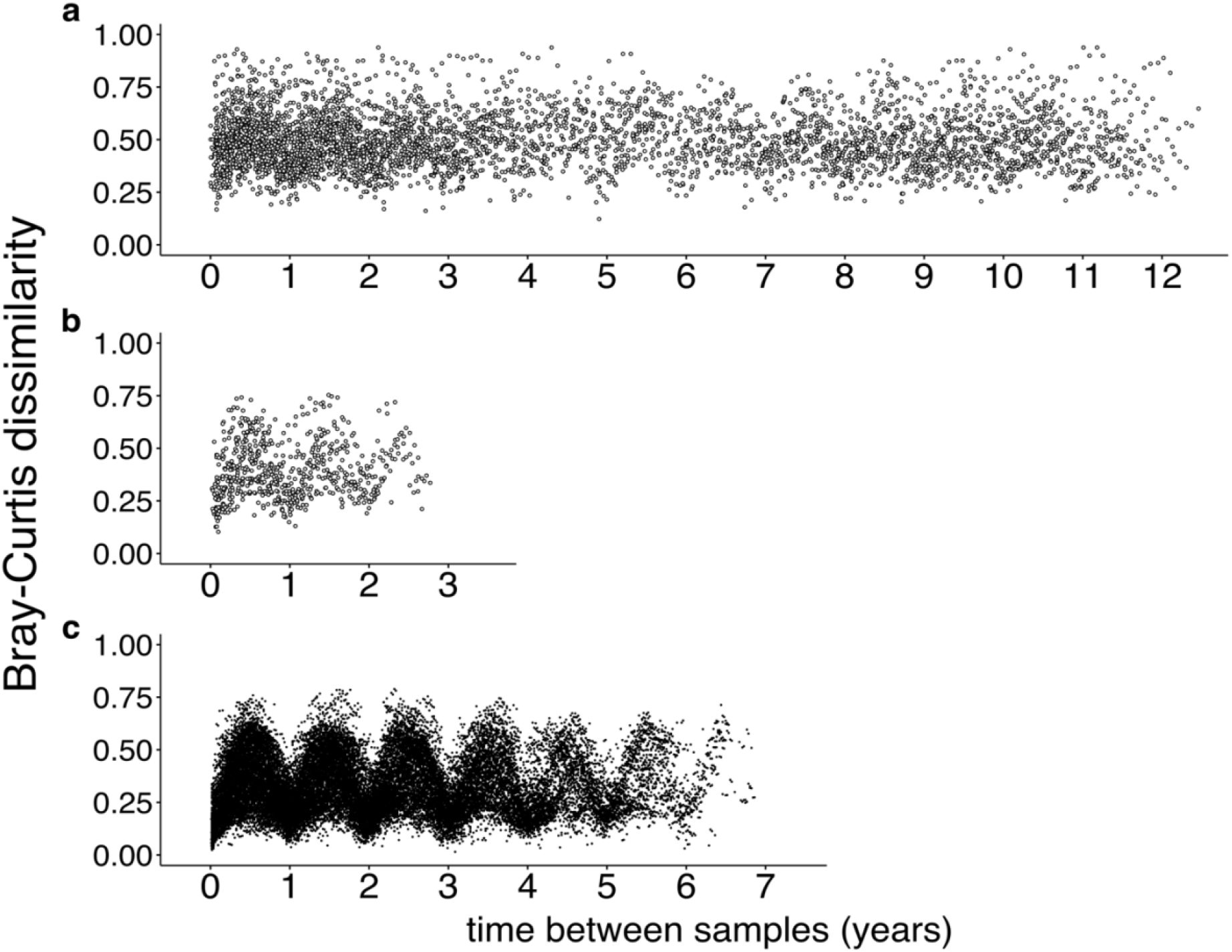
Inter-annual patterns of Bray–Curtis dissimilarity between SAR11 ASV communities. Pairwise SAR11 community dissimilarity was estimated using the ASV rarefied datasets from BATS and WEC. **(a)** Continuous comparison of Bray–Curtis dissimilarities of BATS monthly samples from 1991-1994 and 1996-2002 sequenced using 454 FLX technology. **(b)** Bray–Curtis dissimilarities of BATS monthly samples from 2016 to 2018 sequenced using the Illumina MiSeq platform. **(c)** Bray–Curtis dissimilarities of WEC weekly samples from 2012-2018.

### SAR11 seasonality is configured in an annual two-state pattern

A constrained ordination of the SAR11 communities was generated for BATS and WEC based on rarefied Bray-Curtis dissimilarities to determine whether SAR11 annual periodicity follows a strict seasonal progression. The constrained ordination revealed that the most important factors explaining the variance of the communities are temperature, oxygen and nutrient concentrations (Fig. 4). In both locations, summer and winter SAR11 composition represented the most divergent communities (Fig. 4). Spring and autumn were not characterized by a distinctive set of ASVs but composed mainly of increasing and decreasing contributions of summer and winter SAR11 ASVs, as if these were transition stages between summer and winter clusters. The samples in the constrained ordination forms two clusters that are not determined by astronomical seasons (Fig. 4). Measured environmental variables at BATS and the WEC explained 63% (adjusted 43%) and 60% (adjusted 57%) of the variance in SAR11 communities, respectively. Hierarchical clustering corroborated this pattern in both BATS and WEC data (Figs. S7,S8).

**Figure 4:**
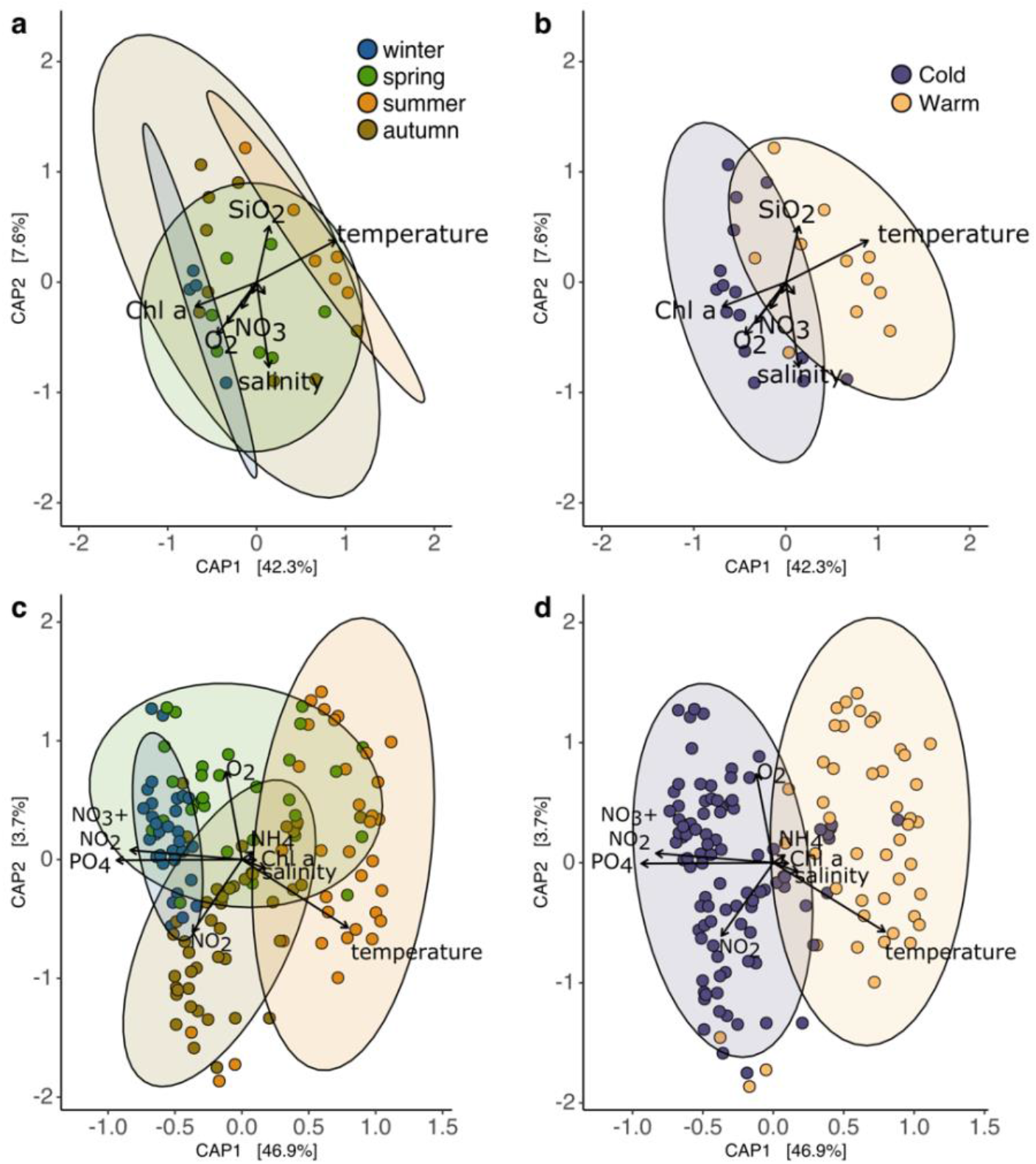
Canonical Analysis of Principle Coordinates (CAP) of SAR11 ASV profiles constrained by physical and chemical environmental parameters. Ordinations were generated with the Bray-Curtis dissimilarities among samples. Bray-Curtis dissimilarities were estimated using rarefied ASV counts. Axis x and y represent the first and second constrained component, respectively. The percentages between brackets are the fraction of the variance explained by each component under the linear model of the explanatory variables. Overlaid arrows represent the explanatory variables, arrow’s length is proportional to the variation explained and its orientation show the direction in which the variable increases. **(a)** BATS monthly samples from 2016 to 2018 color-coded by season. **(b)** BATS monthly samples from 2016 to 2018 color-coded by cold (22 September-1^st^ May) and warm clusters (2^nd^ May – 21 September). **(c)** WEC weekly samples color-coded by season **(d)** WEC weekly samples color-coded by cold and warm clusters (as in panel b).

A linear regression model was used to classify significant periodic ASVs. Two broad seasonal patterns emerged: ASVs with a peak in relative abundance in summer and ASVs peaking in winter. Both seasonal patterns were consistently correlated with the SAR11 ecotype of the ASVs (Fig. 5). Abundant (>0.5%) “warm water” Ia.3 ASVs in both locations, dominate the summer peak pattern. Other ASVs that followed a summer peak pattern at BATS included a single ASV from subclade Ib.2 and multiple low abundance ASVs from subclade IIIa. In the WEC, only one other ASV belonging to ecotype IV showed a summer peak. ASVs that peaked in winter at BATS included those belonging to Ib.2, IIa, IIa.A and IV. These subclades were not represented in the ASVs peaking in winter at the WEC, which instead was comprised of subclades Ia.1, Ib, II, IIa.B and IIb. Thus, while the summer communities at both sites are broadly similar, the ecological niches at both sites in winter select for different SAR11 taxa. Despite having similar abundance peaks, some ASVs had variable peaking times within the season. These may be the result of short-term scale variability within the seasonal progression of SAR11. These results confirm the consistency of SAR11 ecotype seasonality throughout the multiannual surveys and provide evidence supporting a two-state pattern seasonality.

**Figure 5:**
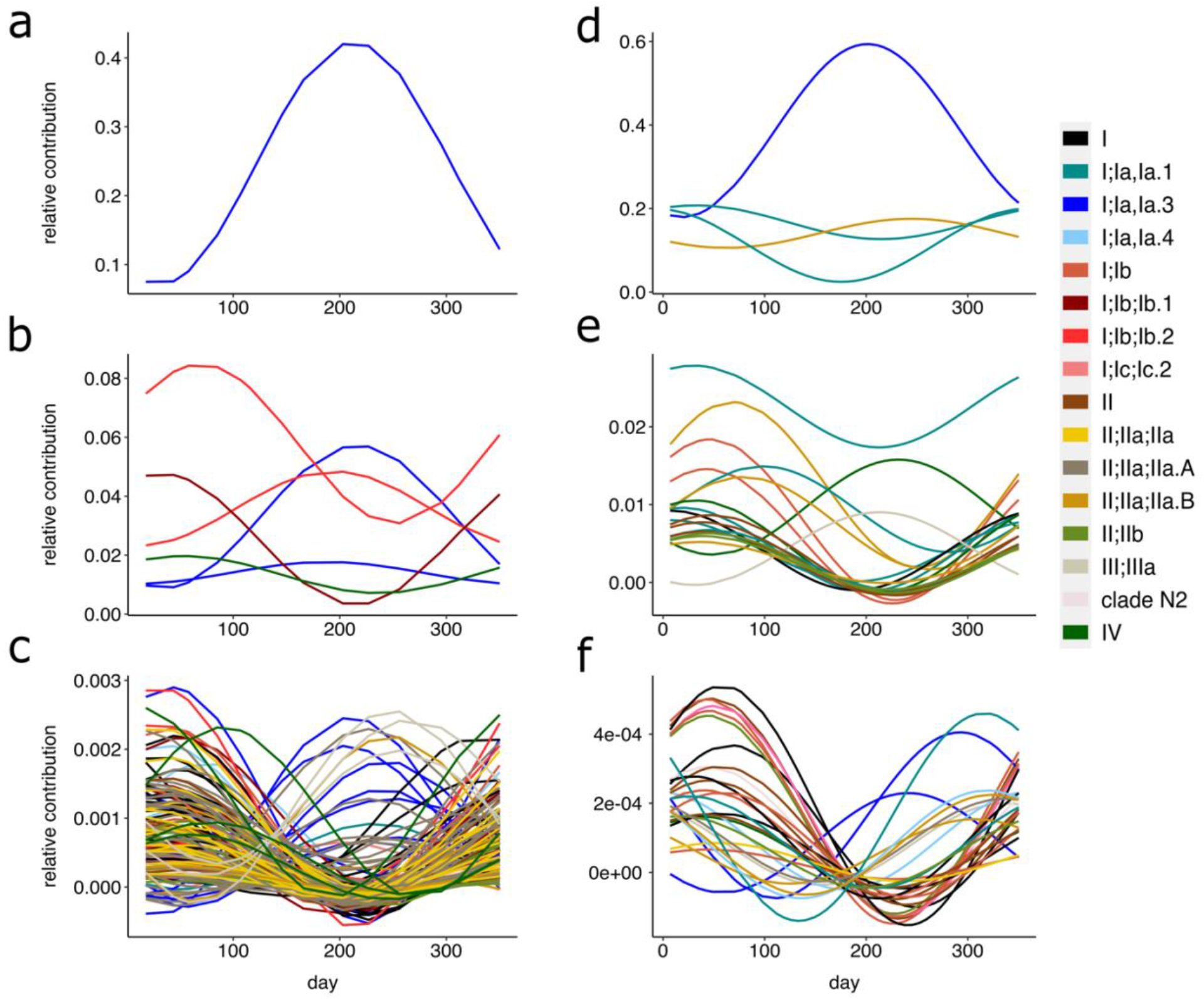
Annual abundance trends of SAR11 ASVs in the Bermuda Atlantic Time-Series and the Western English Channel. A harmonic linear regression model was used to determine significant seasonal trends (p<0.05) of individual ASVs. Two main patterns of ASVs with seasonal contributions peaking in winter and summer were identified. The Bermuda Atlantic Time-Series ASVs are shown in panels **a-c**. **(a)** ASVs with a total relative contribution greater than 5% **(b)** ASVs with a total relative contribution between 0.5% and 5% **(c)** ASVs with a total relative contribution less than 0.5%. The Western English Channel ASVs are shown in panels **d-f**. **(d)** ASVs with a total relative contribution greater than 10% **(e)** ASVs with a total relative contribution between 0.5% and 10% **(f)** ASVs with a total relative contribution less than 0.5%. ASV adjusted model curves are color-coded by ecotype. Day 0 corresponds to the 1^st^ of January.

### Interannual weather variability influences SAR11 short term β diversity and its response to ocean physical and chemical changes

A high-resolution, near complete sampling regime in the WEC from April 2015 to April 2017 (Fig. S9) allowed us to analyse how changes on SAR11 β diversity between weekly subsequent samples are correlated to short-term environmental fluctuations (Fig. 6). We generated consecutive Euclidean distances based on 17 environmental measurements: six weather and eleven physicochemical variables from the ocean surface (Fig. 6a). The Bray-Curtis dissimilarities from consecutive samples showed that during the year 2015, SAR11 communities did not show drastic transitions from week to week (Fig. 6b). In contrast, in 2016 sharp β diversity transitions occurred (Fig. 6b). These steep differentiations were sharpest in the first half of summer and in the transition from autumn to winter. Environmental weekly fluctuations were variable in both years, however the changes in 2016 clearly display similar progressions to those of the SAR11 Bray-Curtis dissimilarities (Fig. 6b). Wavelet coherence analysis corroborated that from the vernal equinox of 2016, Euclidean and Bray-Curtis distances were in phase with a significant oscillation period of 16 weeks. In phase oscillations decoupled after the vernal equinox of 2017 (Fig. S10).

**Figure 6:**
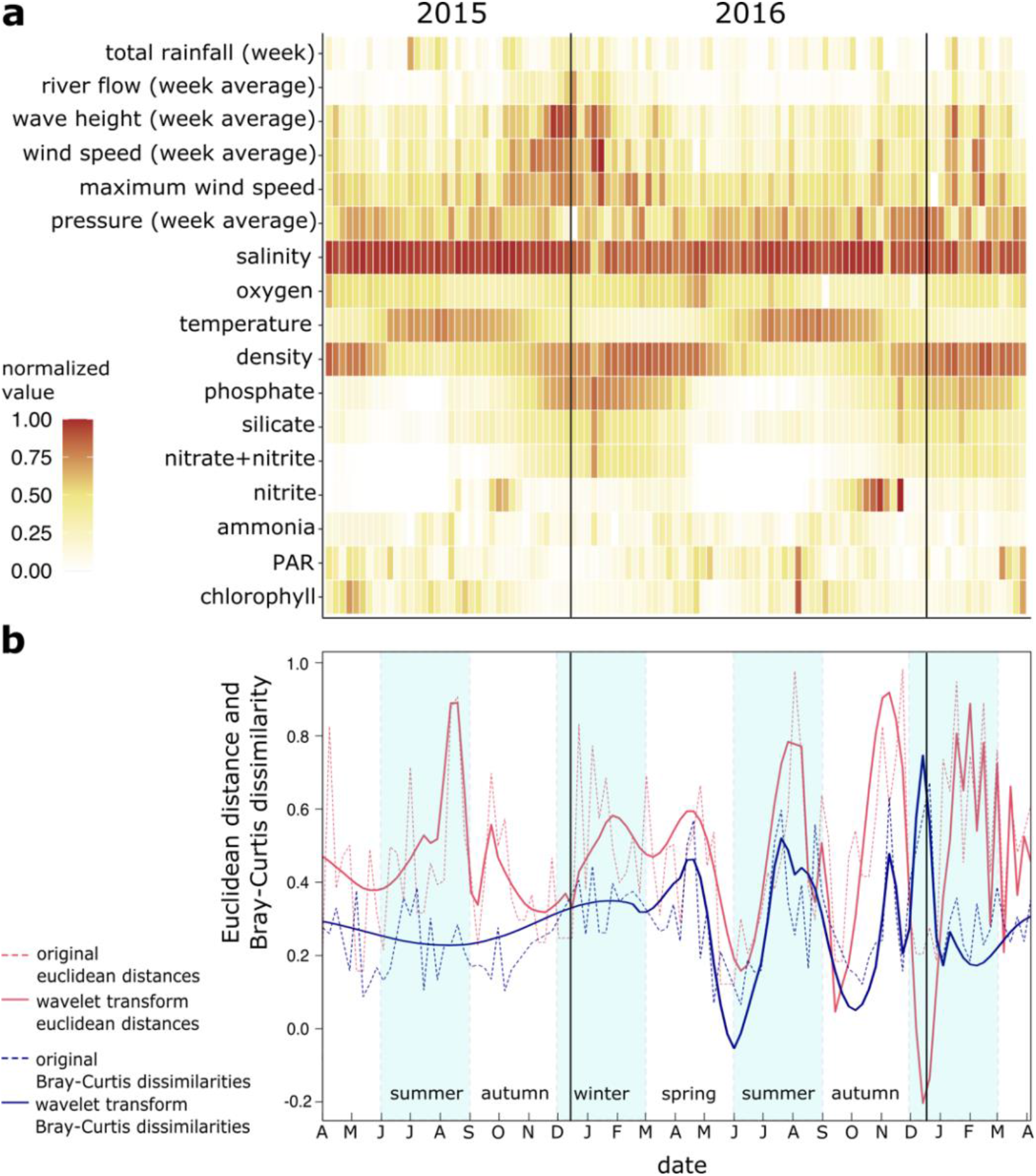
Response of SAR11 community to environmental heterogeneity on a weekly scale at the Western English Channel. **(a)** Normalized values (0 to 1) of the 17 weather and oceanographic environmental parameters used to estimate the Euclidean distance between weekly consecutive samples. **(b)** Wavelet transforms (solid red line) of the weekly consecutive Euclidean distances representing the environmental changes and Bray-Curtis dissimilarities of the SAR11 populations (solid blue line). The original Euclidean distances and Bray-Curtis dissimilarities are shown as dashed lines, red and blue respectively. Panels (a) and (b) are aligned in the x-axis to represent the same week timepoint.

Significantly different environmental variables between the uncoupled and coupled phases (Table S2) were maximum and weekly average wind speed and river flow measurements (p<0.05). Net heat flux, which has previously been shown to drive changes within the whole microbial communities of the WEC (51) were not significantly different between the phases of SAR11 communities (p>0.05; Table S2, Fig. S11). These differences suggest that when unsettled weather conditions occur as in 2015, the surface layer of the ocean (~0-5 m) is well-mixed, homogenizing the physicochemical transitions of the water which in turn causes a gradual seasonal progression of SAR11. Contrarily, in calmer weather conditions, as in 2016, physicochemical seasonal transitions of the surface water are sharper thus reflecting on the short-term response of SAR11 and its progression within the annual cycle.

## Discussion

SAR11 bacteria have been the focus of oceanographic molecular surveys because of their abundance and pivotal role in the ocean biogeochemistry (23,27,29). However, these studies have not been directly comparable due to different sampling periods, sampling frequency, molecular data generation protocols and data analysis. In this study, using a highly defined phylogenetic placement of SAR11 sequences, we analysed for the first time in a comparable manner SAR11 dynamics of two multiannual molecular surveys in the North Atlantic: BATS and WEC.

A major difference between both molecular surveys was the set of primers used, which amplify different 16S rRNA hypervariable regions. Because of the different conservation levels of the 16S rRNA hypervariable between region V4-V5 (WEC) and V1-V2 (BATS) (52–54) (Fig S12), we cannot draw conclusive comparisons regarding the total number of SAR11 ASVs identified, perceived diversity and the proportion of seasonal ASVs predicted in each dataset. Furthermore, the data generated with 454 FLX technology has an output of approximately one order of magnitude less than the 2016-2018 Illumina dataset (0.45 Gb 454FLX vs ~7.5 Gb MiSeq). Nevertheless, by analysing these datasets using the same phylogenetic backbone, we overcame the inherited differences of both time series to provide a panoramic view of SAR11 long-term composition seasonality and the environmental constraints that define it.

Our results showed that SAR11 ecotype seasonality have been consistent in both locations through the studied years. We confirm that temperature and nutrients are strong predictors of SAR11 ecotype geographical distribution and seasonality (3,12,22). However, the specific thresholds that shape SAR11 seasonality and biogeography may be transitional and influenced by other factors such as turbulent mixing, light and biological interactions (viral predation or co-occurring organisms). For example, the warm ecotype Ia.3, thought to be constrained to oligotrophic environments (55), is a seasonal component at WEC, located at latitude 50°N with high nutrient concentrations and temperatures as low as 7 °C. This result may be explained by genomospecies within the ecotype with specific adaptations to different environments (13,56) or by horizontal water transport that disperse planktonic microorganisms stirring its distribution across biogeographical regions (56–58). Whether the Gulf Current constantly seeds the WEC with warm adapted bacteria remains to be investigated. In contrast, the comparison between WEC and BATS suggests that cold ecotype Ia.1 (59), is restricted to a maximum temperature of 20 °C, therefore hardly retrieved from BATS (minimum temperature 19.27 °C). Cultivable representatives, HTCC1062 (Ia.1) and HTCC1072 (Ia.3), have optimal growth temperatures of 16 °C and 23 °C, respectively (60). However, the presence of Ia.3 through a larger range of temperatures than Ia.1 suggests a wider landscape of genetic adaptability or phenotypic plasticity. Ecotypes from clade II also appear to have specific limits of temperature and nutrients shaping their seasonality and occurrence at BATS and WEC. While subclade IIa.A peaked at BATS in winter and early spring, subclades IIa.B and IIb did in the WEC. In contrast, a recent study from the Blanes Bay Microbial Observatory (BBMO), a time series at the coastal oligotrophic NW Mediterranean Sea, showed no seasonal trends of SAR11 clade II ASVs (61). To date, no single cultures of SAR11 II subclades have been reported, limiting our capacity to experimentally verify optimal growth temperatures. The contrasting seasonality of subclade II ecotypes at BBMO, WEC and BATS suggests that environmental factors, i.e. weather conditions or horizontal transport may expand or constrain this ecotype seasonality and distribution.

Weekly SAR11 transitions were uncoupled from the environmental transitions from April 2015 to March 2016 in the WEC. In contrast, from March 2016 to April 2017 transitions were coupled. Significant differences between these years were wind speed and river flow, which we assumed created a more turbulent surface; particularly record breaking weather in winter 2015-2016 (flooding, high winds and temperature) across the United Kingdom (62). We hypothesise that these unsettled weather conditions kept a constant mixing of the surface creating a more homogeneous system and blurring the sharp seasonal transitions that occur in more stable years. Annual differences in wind and river input are common throughout the time series (Fig. S13) suggesting that coupling and uncoupling of SAR11 β diversity to water conditions may be a common phenomenon.

We demonstrate that surface ocean SAR11 populations are influenced by short and long time-scale environmental drivers at two ocean sites that differ dramatically in average productivity and temperature. Patterns of seasonal succession were fundamentally similar between the two sites, but major differences in ecotype dominance were observed that foreshadow changes in ocean ecology likely to occur as ocean temperature increases. Extreme weather events, which are also predicted to increase (63), affect SAR11 surface ecotype progression. Given SAR11 ecotypes utilize different carbon sources and release different gaseous compounds, the result of these changes may have hitherto unknown impacts on the ocean and atmospheric processes.

## Supporting information

see Supplementary methods

A list of samples is provided as Table S1

## Acknowledgements

WEC amplicon sequencing was provided by NU-OMICS. We would like to thank Mark Dasenko, and Oregon State University CGRB [now known as Center for Quantitative Life Sciences (CQLS)] for BATS amplicon sequencing. We would like to thank the crew of the R/V Plymouth Quest, our collaborators at PML for collecting water samples from the WEC, crew of the R/V Atlantic Explorer, and BATS groups for sampling from the BATS site. We thank Shuting Liu and Krista Longnecker for consolidating BIOS-SCOPE metadata. Bioinformatic analyses were conducted using the high-performance computing, ISCA, provided by the University of Exeter.

## Data availability

Amplicon sequence datasets presented in this study have been deposited in NCBI SRA database under the BioProject identifier PRJNA769790 for the Bermuda Atlantic Time-series Study and GeneBank ON706058-ON706222 for the Western English Channel. Metadata, intermediate processing products and the code used in this study are available at https://github.com/lbolanos32/SAR11_BATS_WEC_2022.

