## Supplementary material for "Influence of short and long term processes on SAR11 communities in open ocean and coastal systems": see Supplementary methods

### Supplementary Information

#### Supporting Methods

**Bioinformatics analysis:**

Bermuda Atlantic Time Series

454 FLX (1991-2002) sequence processing

Raw ssf data files were demultiplexed using get.mimarkspackage command from Mothur (1). Fasta and qual files were generated from the individual samples using sffinfo command. Fastq files for each pair of fasta and quality files were generated with the make.fastq. Fastq files were trimmed with CutAdapt (-u 28). Adapter-trimmed sequences were used to generate ASV count table with Dada2 (2). Sequences were quality filtered using the following parameters maxEE=4, rm.phix=TRUE, minLen=190, truncLen= 220. Sequences were denoised to control for homopolymer artifacts. Samples were processed by plates to accurately estimate the error frequency. Taxonomic assignment was performed with the assignTaxonomy command and the silva non redundant database V123 (3). Finally, an ASV table was generated for each plate. ASV tables were merged using merge_phyloseq command in Phyloseq R package (4).

Illumina MiSeq (2015-2018) sequence processing.

Primers from the demultiplexed raw paired-end fastq files were cut using the CutAdapt (5), removing 20 bases from forward files and 18 from reverse files that matched the 27F and 338 RPL primer lengths, respectively. Sequences were quality filtered using the following parameters maxEE=(2,2), truncQ=2, minLen=190, truncLen= 220,190, maxN=0. Quality filtered sequences were dereplicated and merged with dada2 R package, version 1.2 (2). Sequences from the same run were processed together to accurately estimate the error frequency. Taxonomic assignment was performed with the assignTaxonomy command and the silva non redundant database V123 (3). ASV tables were generated for each plate. ASV tables were merged using merge_phyloseq command in Phyloseq R package (4).

SAR11 ASVs taxonomic assignment

SAR11 ASV tables were obtained by sub setting the sequences assigned to the order "SAR11_clade". SAR11 sequences from the 454 FLX and MiSeq datasets were aligned using mafft (Katoh et al., 2002). The alignment were manually trimmed and the resulting file was collapsed to dereplicate potential duplicated ASVs generated after trimming. The ASV counts were collapsed accordingly. Finally, a fasta file of the unique ASVs were generated. SAR11 sequences were placed in a curated reference tree (6) using Phyloassigner version v6.166 (7). The result of the highly-resolved phylogenetic placement was used to update the taxonomic file.

Western Channel Observatory (2012-2018)

Illumina MiSeq sequence processing.

Primers from the demultiplexed raw paired-end fastq files were cut using the CutAdapt (5), removing the primer sequences based in all potential orientations. Sequences were quality filtered using the following parameters maxEE=(2,2), truncQ=2, truncLen= (240,200), maxN=0. Quality filtered sequences were dereplicated and merged with dada2 R package, version 1.2 (2). Sequences from the same run were processed together to accurately estimate the error frequency. Taxonomic assignment was performed with the assignTaxonomy command and the silva non redundant database V138 (3). ASV tables were merged using merge_phyloseq command in Phyloseq R package (4).

SAR11 ASVs taxonomic assignment

SAR11 ASV tables were obtained by sub setting the sequences assigned to the order "SAR11_clade". The SAR11 ASVs were placed in a curated reference tree (6) using Phyloassigner version v6.166 (7). The result of the highly-resolved phylogenetic placement was used to generate an updated taxonomic file.

**Temporal diversity and influence of environmental variables on SAR11 composition:**

A wavelet coherence analysis was performed on the weekly WEC sampling from April 2015 to April 2017 to determine whether betta diversity differences mirror the environmental changes (R package WaveletComp (8)). WEC time-series is regularly sampled on Mondays, unless it is

rescheduled or cancelled due to weather conditions. To achieve fixed equidistant sampling dates (every 7 days), we reassigned the date to the closest Monday and linearly interpolated the missing data.
The data of 17 environmental variables were standardized to values between 0 and 1, based on the minimum and maximum values of each, using the formula: X'=x-min(x)/max(x)-min(x). These standardized values were used to generate Euclidean distances between subsequent samples representing the change in the environment between weeks. Beta diversity used throughout the different analysis of this manuscript was generated based on rarefied amplicon counts (1000 reads). A coherence analysis between Bray-Curtis dissimilarities and Euclidean distances of the environment was performed to explore the potential timeframes where both time-series are correlated.

### Supporting Results

#### SAR11 ecotype seasonality is persistent through multiannual time-series in the Sargasso Sea and Western English Channel

SAR11 fraction was comprised of 557 ASVs (454 FLX) and 697 (MiSeq) at BATS and 168 at WEC before filtering by minimum abundance (>.0015). From these, 518, 650 and 147 were assigned to terminal nodes of the SAR11 phylogenetic tree, respectively (Fig S3). For this study, we considered the full internal and terminal nodes collection of SAR11 ASVs.

We evaluated whether the SAR11 ASVs from the BATS period of 2016-2018 with relative abundance > 0.05% presented a significant periodicity (Fisher *G*-test, $p < 0.05$). From the top 169 ASVs, 49.7% displayed a significant periodicity through the analysed years (representing 65.02% of the total reads). From the 11 most abundant ASVs (>2%), five (45.45%) showed a significant periodicity (Fig. S1). Similarly, we evaluated the periodicity of SAR11 ASVs retrieved at the WEC. We analysed the total 144 WEC ASVs, from which 88.2% were determined to display significant periodicity (representing 99.4% of the total reads), including the 10 most abundant ASVs (>2%) (Fig. S2). SAR11 experiencing seasonal periodicity is much higher in WEC than BATS, most likely due to the much bigger seasonal shifts or the V4-V5 region higher conservation (Fig. S12) masking minority ASVs dynamics.

#### Decreasing primary productivity is negatively correlated with the increasing temperature observed in open ocean, but not coastal site.

BATS is located in the western North Atlantic subtropical gyre (31°40′N,64°10′W). At this open ocean site, the annual cycle of the water column is characterized by a strong vertical thermal stratification in summer and the deepening of the mixed layer in winter (150 to 300m) (9). In the years covered by this study, surface water temperature ranged from 19.27° C to 29.58° C (Fig.

S4). Nitrogen and phosphate nutrients were at or close to analytical detection limits (30 nmol $kg^{-1}$ before 2004 and 1 nmol $kg^{-1}$ after 2004 (10) throughout all the sampling, reflecting the oligotrophic nature of the gyre. Chlorophyll concentration showed a clear peak in the winter-spring transition. Over the course of the time-series, spanning 27 years, the peak chlorophyll concentration at 5m decreased while minimum annual temperatures have increased (Figs. S1,S3). The L4 coastal station (50°15'N,4°13'W) is located in the Western English Channel. During the seven years analysed in this study, surface water temperature ranged between 7.62° C to 19.16° C (Fig. S5). Nitrate and nitrite, silicate, and phosphate concentrations peaked in winter and were depleted during summer. Ammonium concentrations varied throughout the year. Chlorophyll concentration displayed multiple annual peaks, with the highest occurring in spring and not showing a correlation with long-term minimum temperature changes as in BATS (Fig. S5, S6).

**SAR11 community seasonality is consistent with a two-state pattern (warm-cold) at WEC and BATS**

Hierarchical clustering corroborated a two-state pattern (warm-cold) at WEC and BATS in both, BATS and WEC (Fig. S7, S8). Two major clusters were described corresponding to summer and winter SAR11 communities, while spring and autumn communities were positioned depending on whether they were more closely related to summer or winter.

### Supplementary Tables and Figures

**Supplementary Table 1. Time series samples used in this study and their corresponding environmental data (Provided as an independent document).**

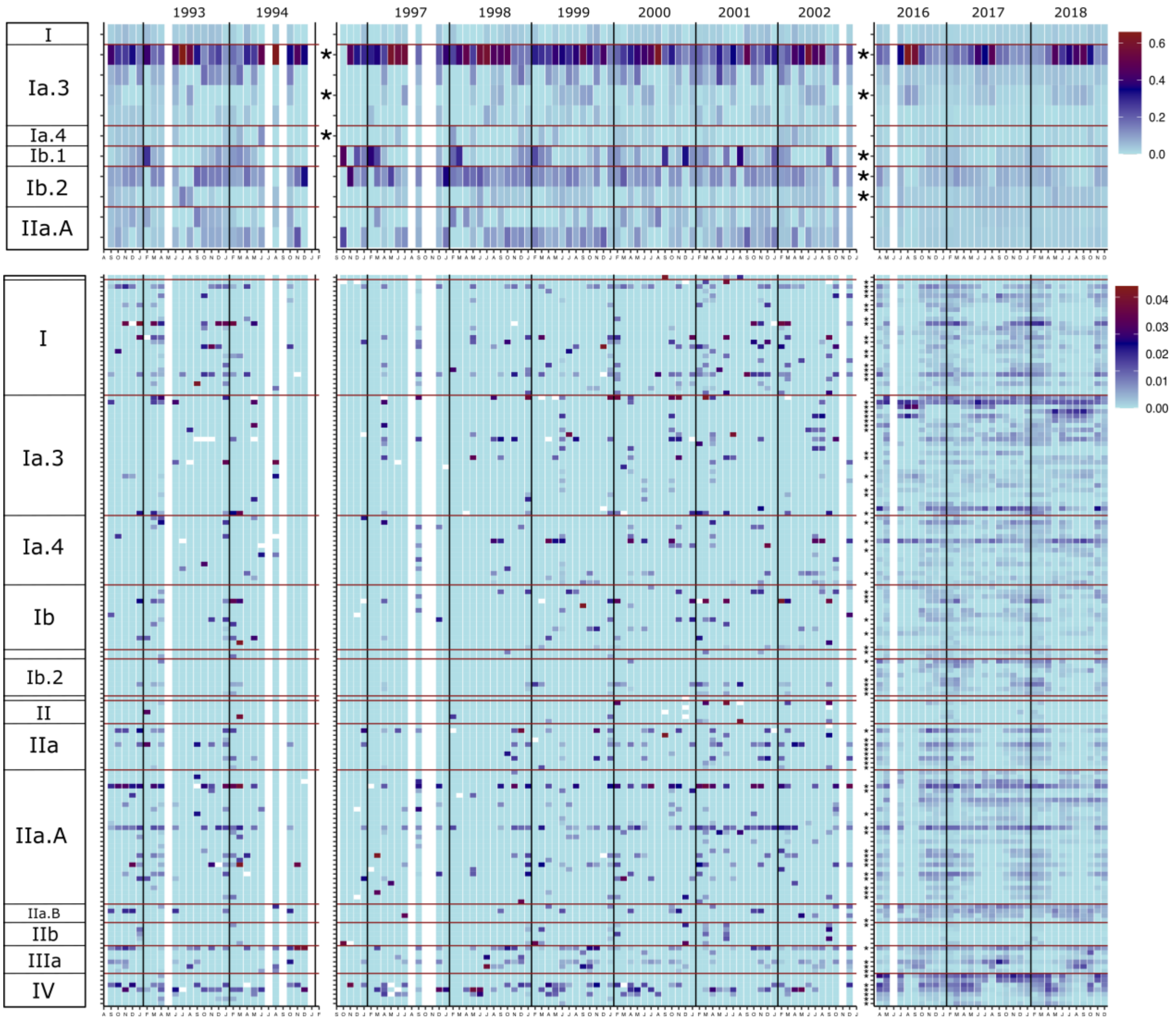


**Supplementary Figure 1. Heat map depicting the monthly relative abundance of the top 169 SAR11 ASVs from the Bermuda Atlantic Time-series Study.** ASVs with a relative abundance > 0.05% were selected. The ASVs are divided into two groups based on its contribution. The top panel displays ASVs with a relative contribution greater than 2% (top 11 ASVs). The bottom panel displays ASVs with a relative contribution less than 2% (158 ASVs). The ASVs are sorted in the y-axis by ecotype for both panels. Within ecotypes, ASVs are sorted vertically by decreasing relative contribution to the dataset. Asterisk in the y-axis indicates that the ASV displays a significant periodic pattern based on a Fisher G-test ($p < 0.05$) (See Supplementary Materials and Methods). Fisher G-test were done independently for the period between 1991-2002 and 2016-2018.

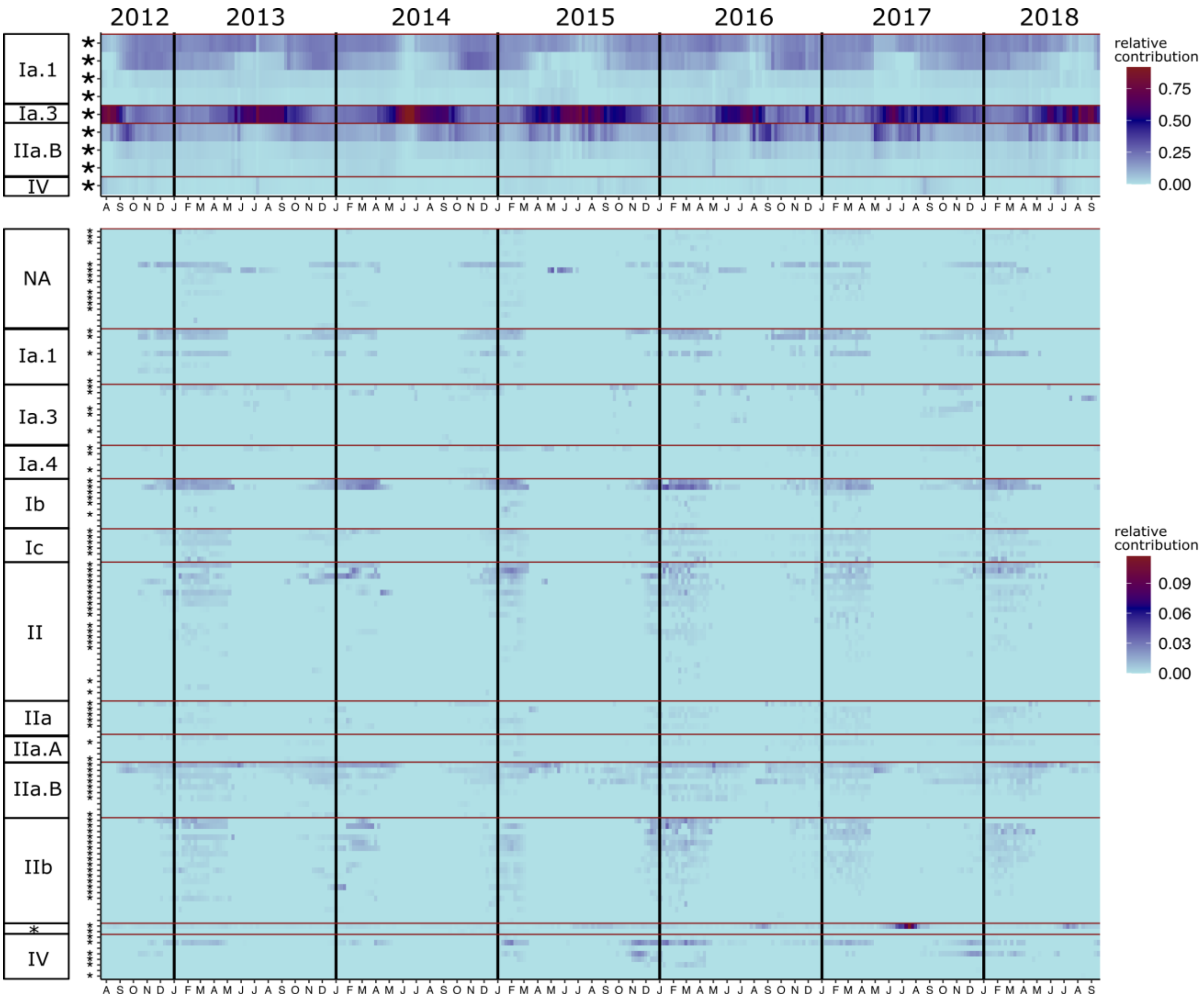


**Supplementary Figure 2. Heat map depicting the weekly relative abundance of the SAR11 ASVs from the Western English Channel.** All SAR11 ASVs (144) are depicted. Samples with less than 1000 SAR11 reads were removed. The missing weekly samples were interpolated using the final dataset. The ASVs are divided into two groups based on its contribution. The top panel displays ASVs with a relative contribution greater than 2% (top 9 ASVs). The bottom panel displays ASVs with a relative contribution less than 2% (158 ASVs). ASVs are sorted in the y-axis by ecotype for both panels. Within ecotypes, ASVs are sorted vertically by decreasing relative contribution to the dataset. Asterisk in the y-axis indicates that the ASV displays a significant periodic pattern (estimated in between $1^{st}$ September 2013 and $28^{th}$ August 2018) based on a Fisher G-test ($p < 0.05$) (See Supplementary Materials and Methods). [$^{X}$] = clade N2.

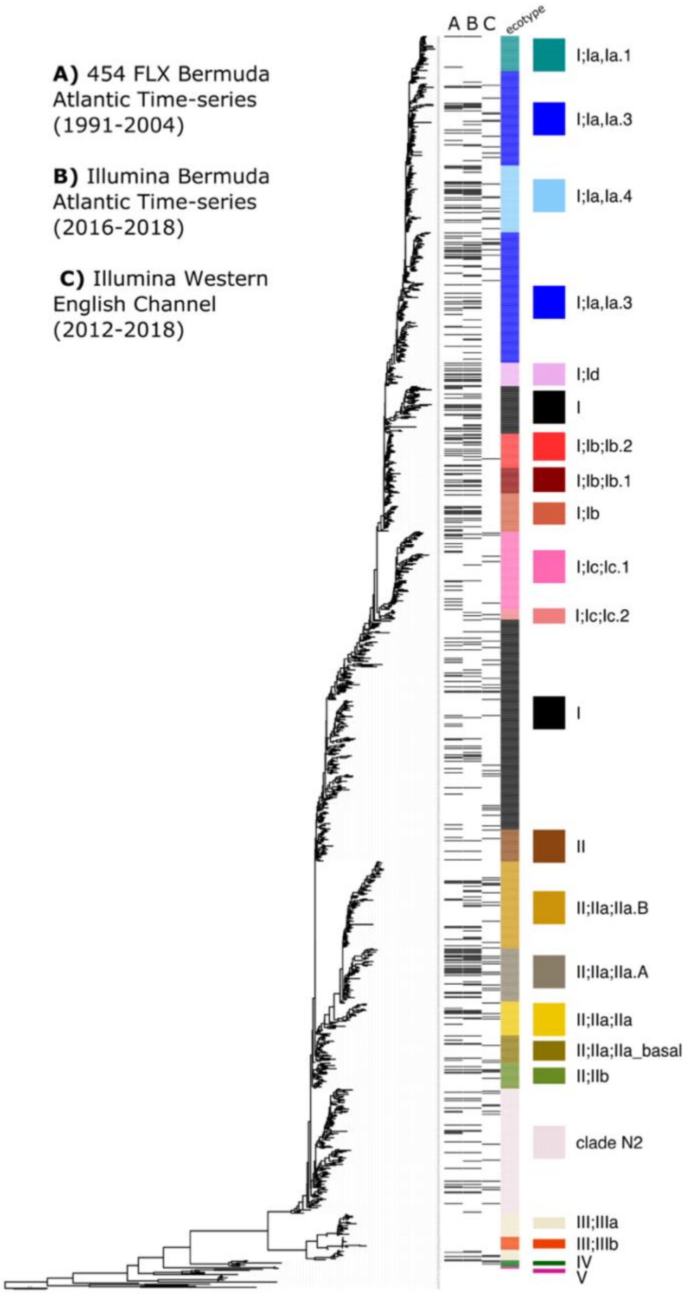


**Supplementary Figure 3. Terminal node ASV profiles for the different datasets: BATS 454FLX, BATS MiSeq and WEC MiSeq.** The full-length 16S rRNA phylogenetic tree used as database to assign the ASV sequences is depicted on the left (Bolaños et al., 2021). Three columns showing the presence (black) and absence (white) of ASVs for each terminal node are shown in front of the corresponding tip of the tree. Each column corresponds to a dataset (**A:** BATS 454FLX**, B:** BATS MiSeq and **C:** WEC MiSeq). The fourth column on the right side show a color-coded relational table for each tip on the tree to the ecotype it belongs. Ia.1 WEC sequences were placed in internal nodes due to its high conservation and therefore not shown in this figure.

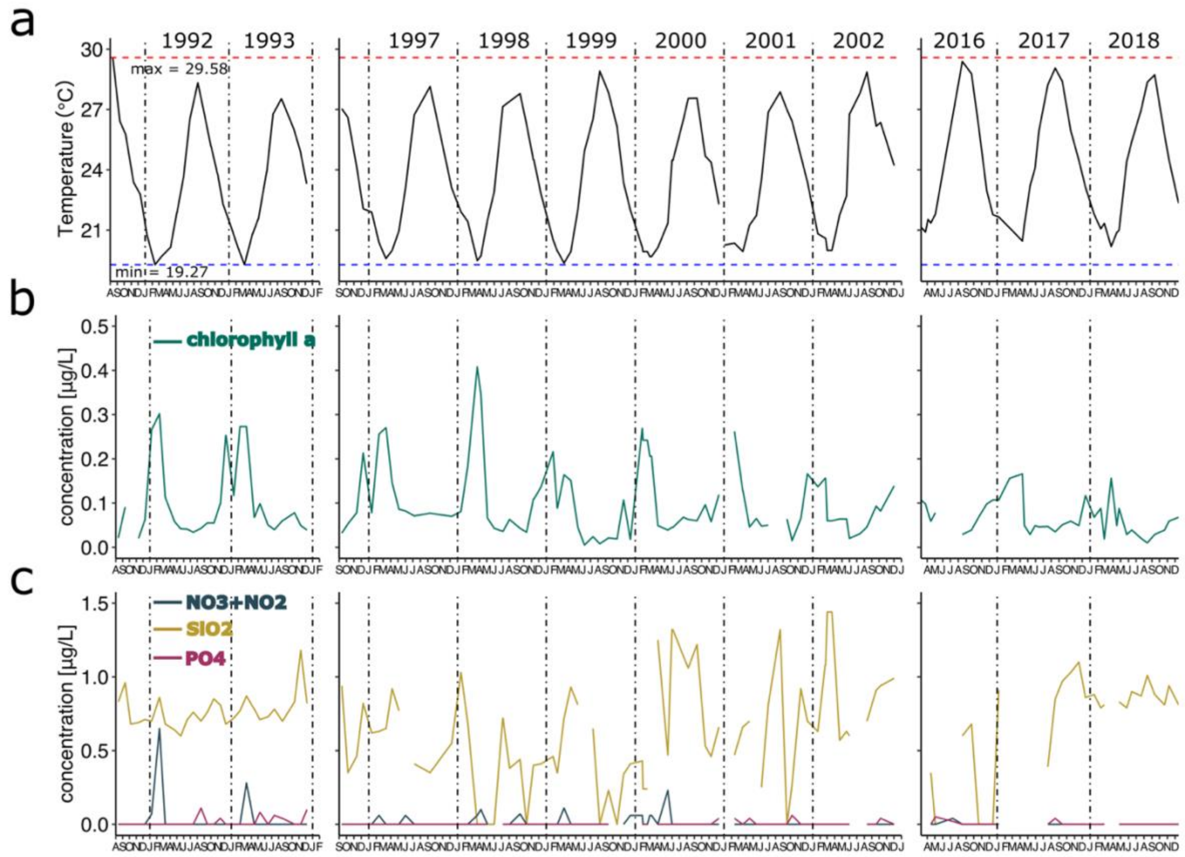


**Supplementary Figure 4. Surface (5m) physicochemical environmental variables measured in the Bermuda Atlantic Time-series Study from 1991 to 2018.** The discontinuity of the lines represents missing values. **(a) Temperature**. The maximum temperature from the measured time ranges is shown as a red dashed line. The minimum temperature is shown as a blue dashed line. **(b) Chlorophyll a**. Low chlorophyll concentrations (<0.25 ug/L) were present for most of the time-range, except for early and late 90s winter to spring transitions. A long-term decreasing trend in chlorophyll a concentration from the early 90s to late 2010s is evident **(c) Nutrients.** Nitrate and nitrite (blue) and phosphate (pink) were close to the detection limits in most of the samples. Silica (yellow) displayed variable concentrations throughout the sampled years without a distinguishable pattern.

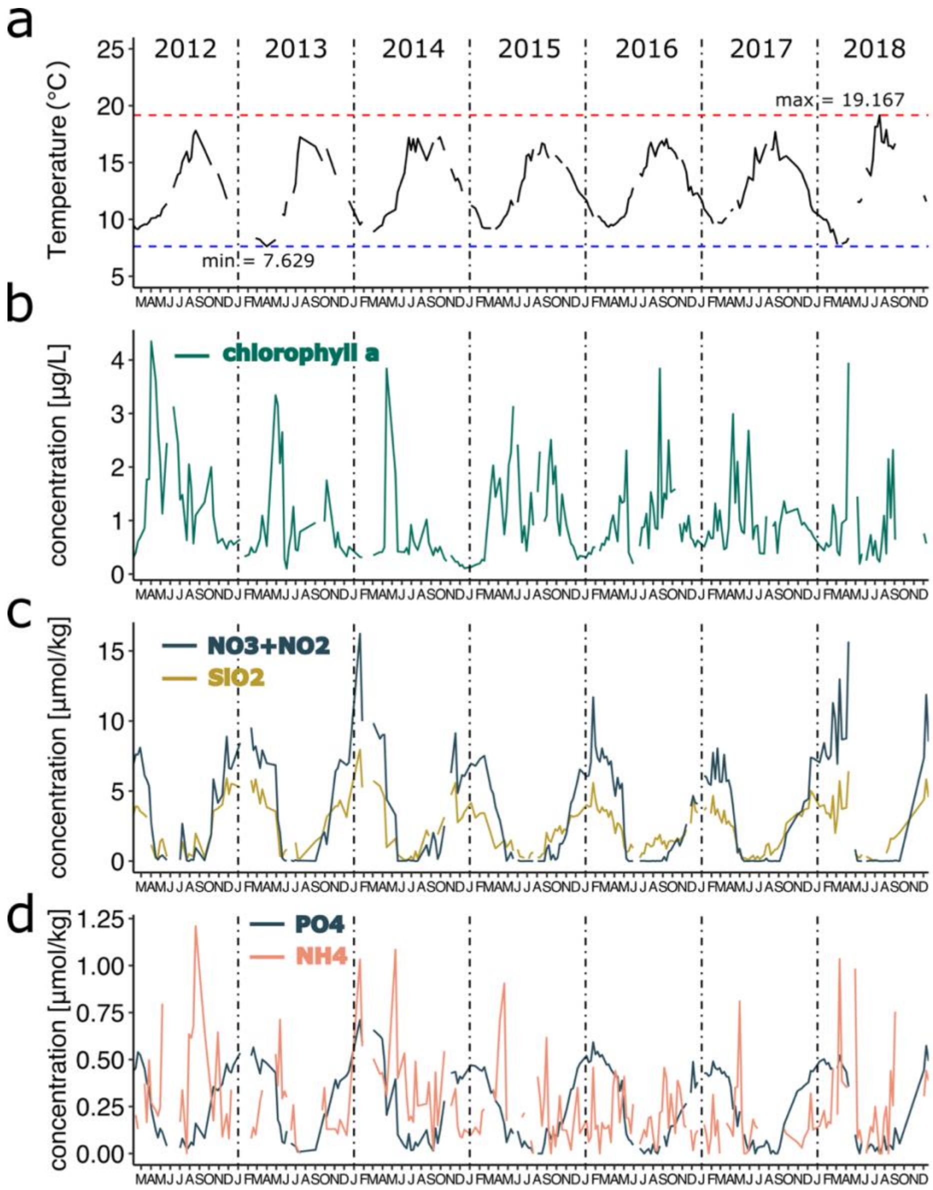


**Supplementary Figure 5. Surface (5m) physicochemical environmental variables measured in the Western English Channel (station L4) from 2012 to 2018.** The discontinuity of the lines represents missing values. **(a) Temperature**. The maximum temperature from the measured time ranges is shown as a red dashed line. The minimum temperature is shown as a blue dashed line. **(b) Chlorophyll a concentration.** High concentration peaks display inter-annual variability; however, spring and autumn bloom peaks are consistent throughout the sampled years. **(c) Nutrients.** Nitrate and nitrite (blue) and silica (yellow) displayed a seasonal oscillation with high concentrations in autumn and winter, while being close to detection limits in spring and summer. **(d) Nutrients**. Phosphate (blue) in panel (d) displayed a seasonal oscillation similar to Nitrate and nitrite (blue) and silica (gold) in panel (c). Ammonium concentrations were highly variable within the range of 0 to 1.25 umol/Kg.

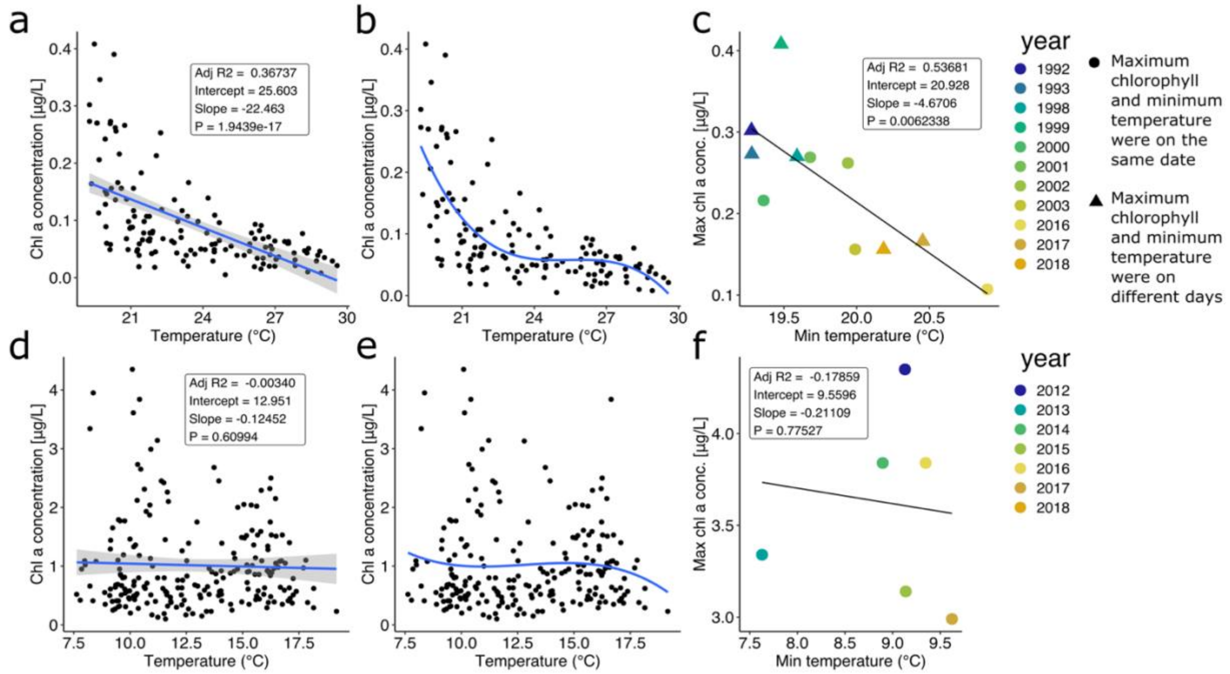


**Supplementary Figure 6. Correlation of chlorophyll a concentrations and temperature of BATS and WEC time-series. (a)** BATS linear regression of concurrent measurements of chlorophyll and temperature. **(b)** BATS polynomial regression of concurrent measurements of chlorophyll and temperature. **(c)** BATS linear regression of the maximum chlorophyll concentration and the minimum temperature for each year sampled during the time-series. **(d)** WEC linear regression of concurrent measurements of chlorophyll and temperature. **(e)** WEC polynomial regression of concurrent measurements of chlorophyll and temperature. **(f)** WEC linear regression of the maximum chlorophyll concentration and the minimum temperature for each year sampled during the time-series. In **(c)** and **(f)** shape represent whether the maximum chlorophyll concentration and minimum temperature measurements belong from the same sampling day (circle) or in a different day (triangle).

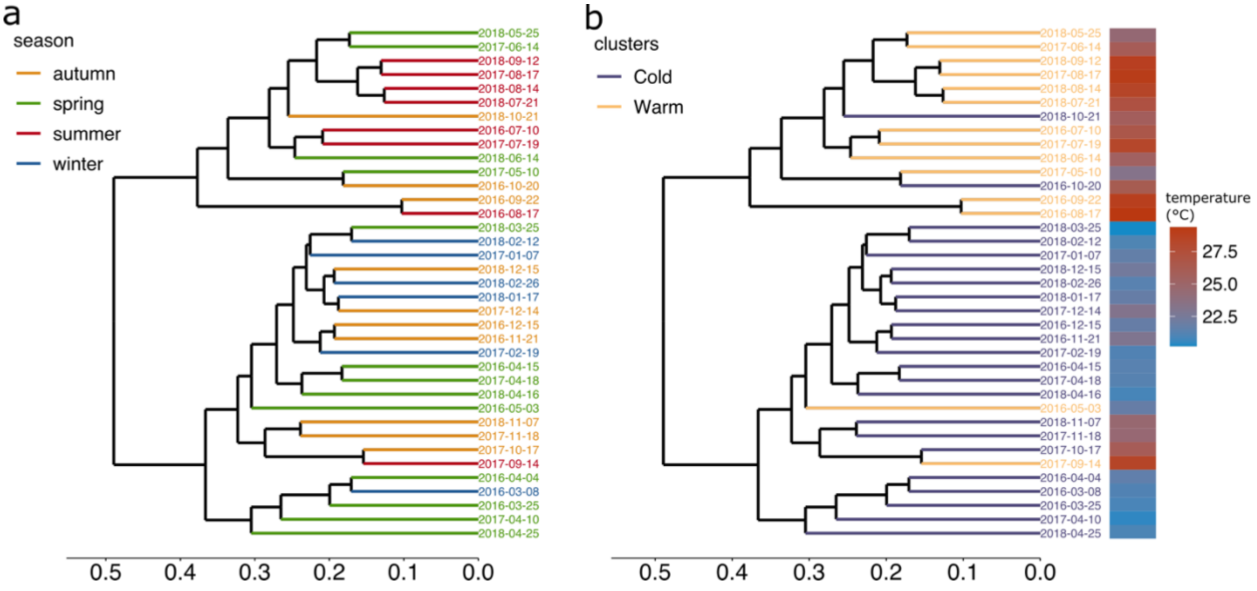


**Supplementary Figure 7. ASVs dendrograms depicting the hierarchical clustering of the samples based on the SAR11 ASV profiles at the Bermuda Atlantic Time-series Study (2016-2018). (a)** ASVs dendrogram defined by hierarchical clustering of the SAR11 community at 5 m depth color coded by seasons in the northern hemisphere. **(b)** ASVs dendrogram defined by hierarchical clustering of the SAR11 community at 5 m depth color coded by a broader definition of seasonality for SAR11 based on the dynamics presented in figures 4 and 5. "Cold" comprises samples from 22 September to 1st May and "Warm" from 2nd May to 21 September. Heatmap (right) represents the temperature recorded for the sample on the corresponding node for both dendrograms.

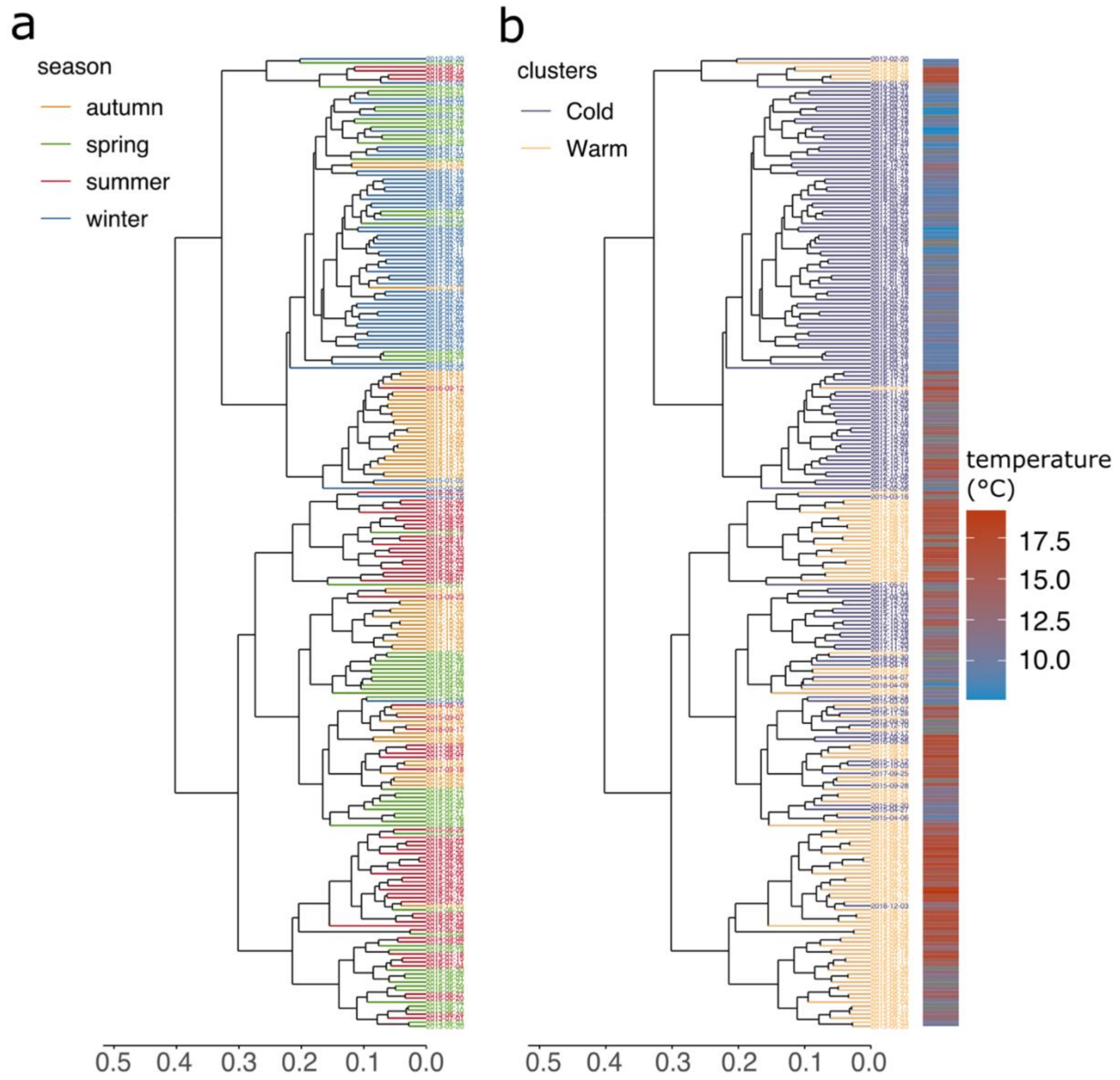


**Supplementary Figure 8. ASVs dendrogram depicting the hierarchical clustering of the samples based on the SAR11 ASV profiles at the Western English Channel (2012-2018). (a)** ASVs dendrogram defined by hierarchical clustering of the SAR11 community at 5 m depth color coded by seasons in the northern hemisphere. **(b)** ASVs dendrogram defined by hierarchical clustering of the SAR11 community at 5 m depth color coded by a broader definition of seasonality for SAR11 based on the dynamics presented in figures 4 and 5. “Cold” comprises samples from 22 September to 1st May and “Warm” from 2nd May to 21 September. Heatmap (right) represents the temperature recorded for the sample on the corresponding node for both dendrograms.

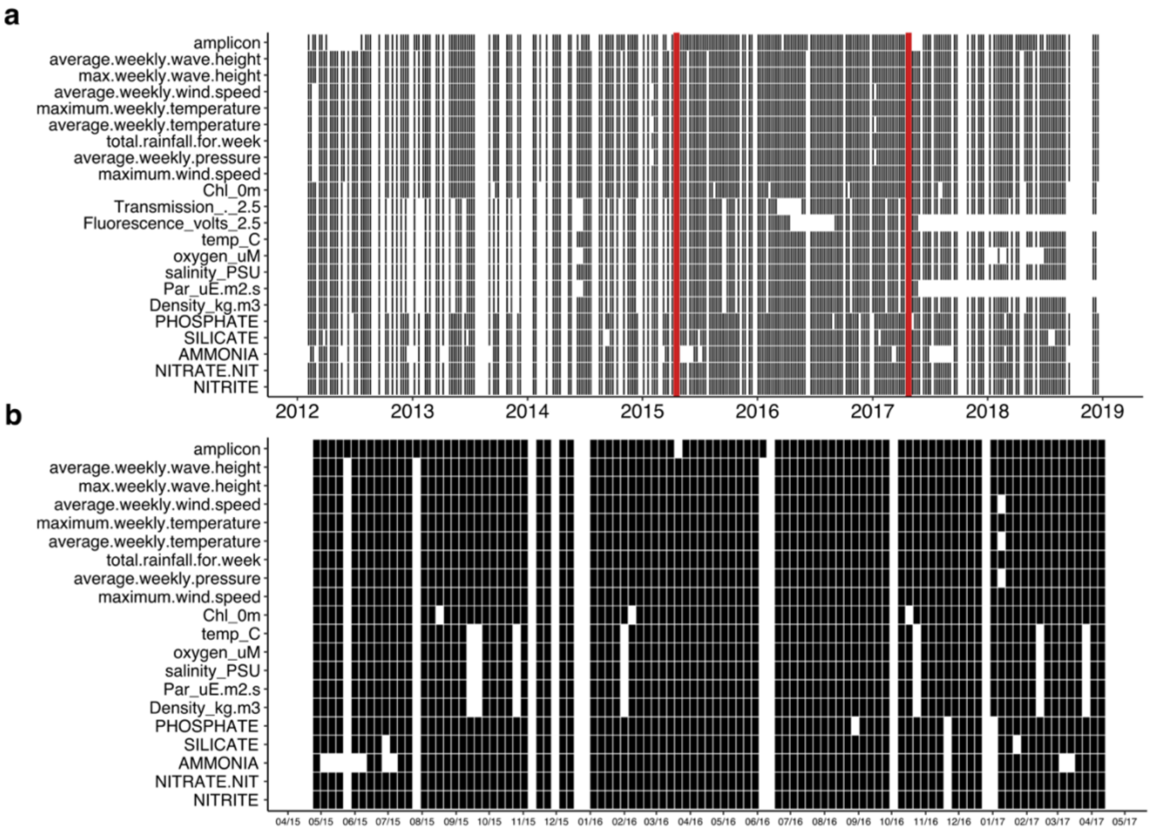


**Supplementary Figure 9. Binary representation of the presence and absence of amplicon datasets and their corresponding environmental measurements at the Western English Channel from 2012 to 2018.** Black squares represent the presence of the indicated measurement, while whites represent the absence. **(a)** Representation of the collected measurements throughout the complete time-series analysed in this study **(b)** Fraction of the measurements taken between April 2015 and April 2017, indicated between red lines in panel (a).

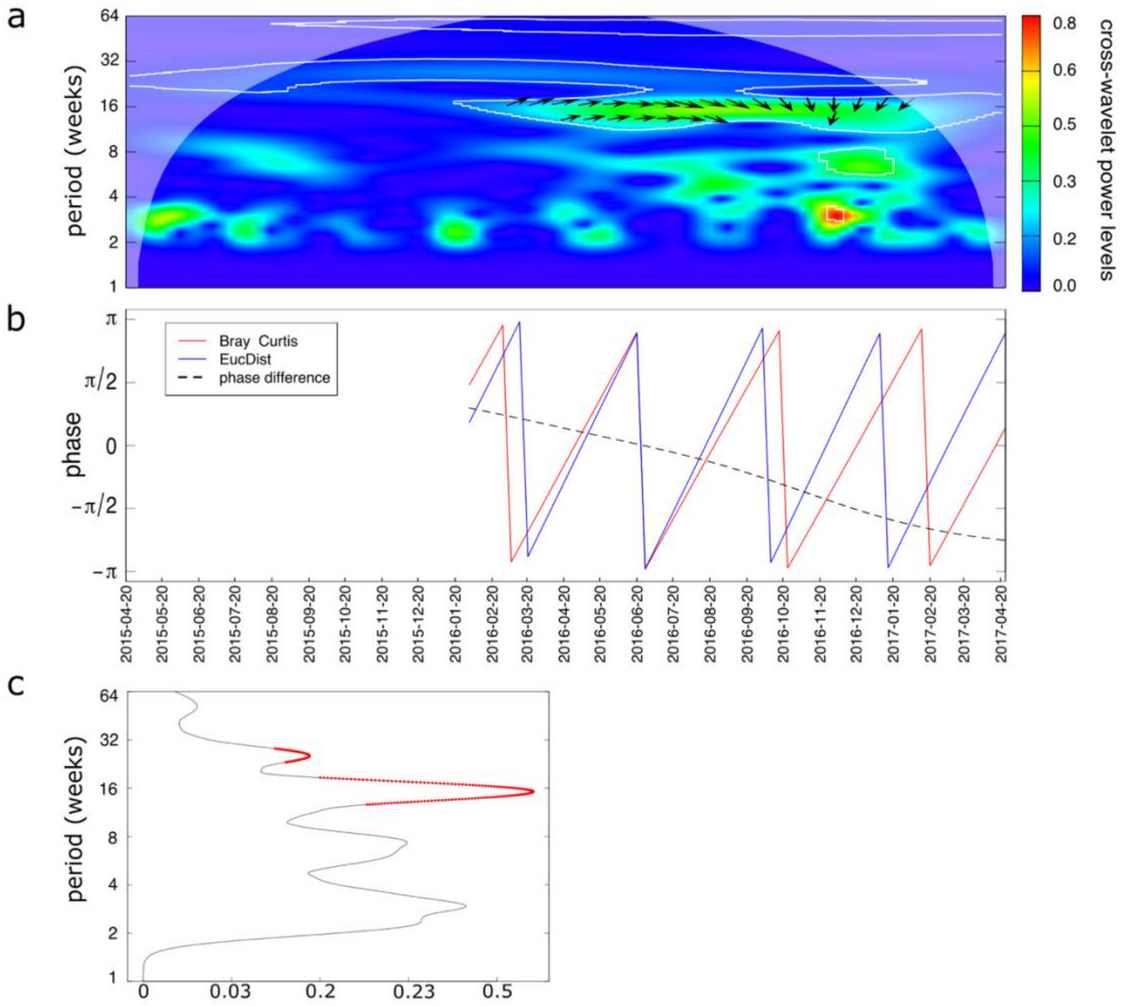


**Supplementary Figure 10. Wavelet coherence analysis between Bray–Curtis dissimilarity and Euclidean distances in the period between April 2015 and April 2017 at the Western English Channel. (a)** Cross-wavelet power levels represented as a color gradient. Periods with significant coherence (P > 0.05) are illustrated by a white line. The angle of the arrows indicate whether the two variables are in phase (positive correlation) and its direction if these are lagged. In phase with x followed by y corresponds to a 45 degree angle (↗), out of phase with y followed by x corresponds to a 135 degree angle (↘). In phase and out of phase without lagging periods are represented by → and ←, respectively. **(b)** In phase representation of the two variables (red and blue) overlapped by the phase difference progression (dashed black line). **(c)** Average coherence for the significant in phase period (P > 0.05) is indicated in red.

| Welch two Sample t-test (2015/2016 vs 2016/2017) | | | |
|---|---|---|---|
| variable tested (H0: μ1=μ2) | t | df | p-value |
| maximum wind speed | -2.352 | 88.447 | 0.02089* |
| average weekly river flow | -2.1961 | 62.843 | 0.02514* |
| average weekly wind speed | -2.1718 | 65.978 | 0.0316* |
| average weekly wave height | 1.9793 | 73.84 | 0.05151 |
| Photosynthetic Active Radiation (PAR) | 1.9609 | 85.135 | 0.05316 |
| nitrite | 1.9121 | 89.534 | 0.05906 |
| Net Heat Flux (NHF) | -1.669 | 714.49 | 0.09555 |
| salinity | -1.6604 | 78.108 | 0.1008 |
| total rainfall (week) | -1.4042 | 76.957 | 0.1643 |
| average weekly pressure | 1.1376 | 89.361 | 0.2583 |
| oxygen | -0.9484 | 83.873 | 0.3457 |
| temperature | -0.53653 | 82.727 | 0.593 |
| phosphate | 0.40487 | 80.547 | 0.6866 |
| nitrate+nitrite | 0.27455 | 77.972 | 0.7844 |
| chlorophyll | -0.17034 | 80.85 | 0.8652 |
| ammonia | -0.1275 | 56.203 | 0.899 |
| density | 0.088802 | 83.463 | 0.9295 |
| silicate | -0.024663 | 74.383 | 0.9804 |

**Supplementary Table 2. Welch's two sample t-test results comparing coupled and uncoupled measurements between April 2015 and April 2017 at the Western English Channel.** * $p\text{-value} < 0.05$.

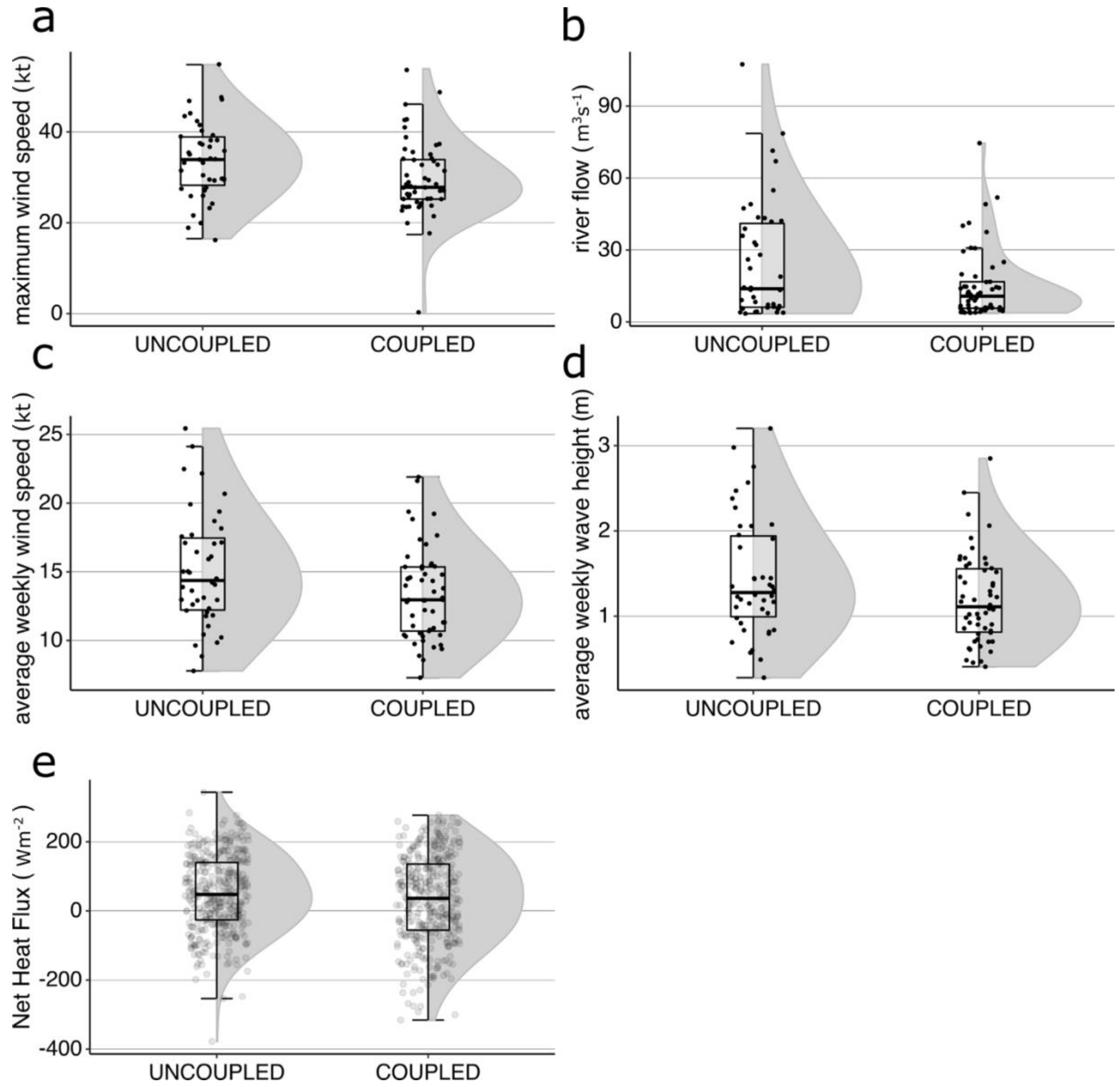


**Supplementary Figure 11. Violin plot overlapped by a boxplot and the corresponding measurements as black points of the variables that showed a significant difference (a,b,c) during the uncoupled and coupled periods from April 2015 to April 2017 at the Western English Channel. Average weekly wave height (d) and Net Heat Flux (e) are also shown for comparison.**

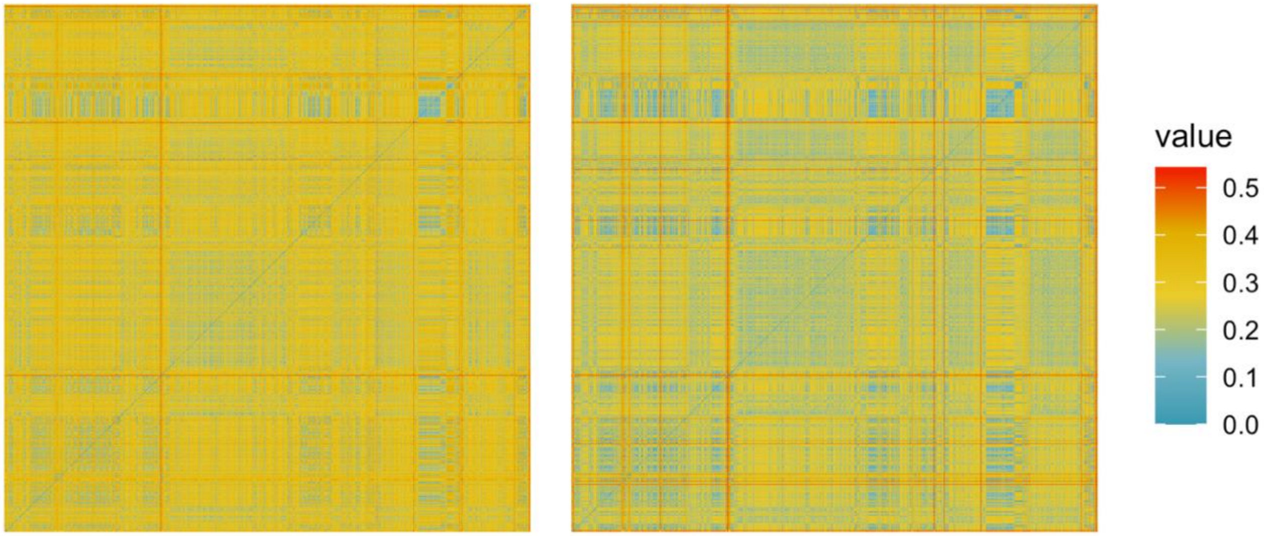


**Supplementary Figure 12. Heatmap matrix depicting the pairwise genetic distance of the V1-V2 and V4-V5 hypervariable regions between the full length sequences used to reconstruct the SAR11 phylogenetic database (Fig S3, Bolaños et al., 2021).** Genetic distances were estimated on the fragments excised from the original alignment using dist.alignment command from the seqinr (Charif D, Lobry J 2007) R package. V1-V2 region is shown on the left panel and V4-V5 on the right. The color-code ranges from identical sequences (0 distance, blue) to 50% identity (0.5 distance, red). The pairwise distances of V1-V2 and V4-V5 are significantly different (Welsch t = 532.88, df = 2683441, p-value < 2.2e-16), supporting that the level of conservation of V4-V5 is higher than the V1-V2.

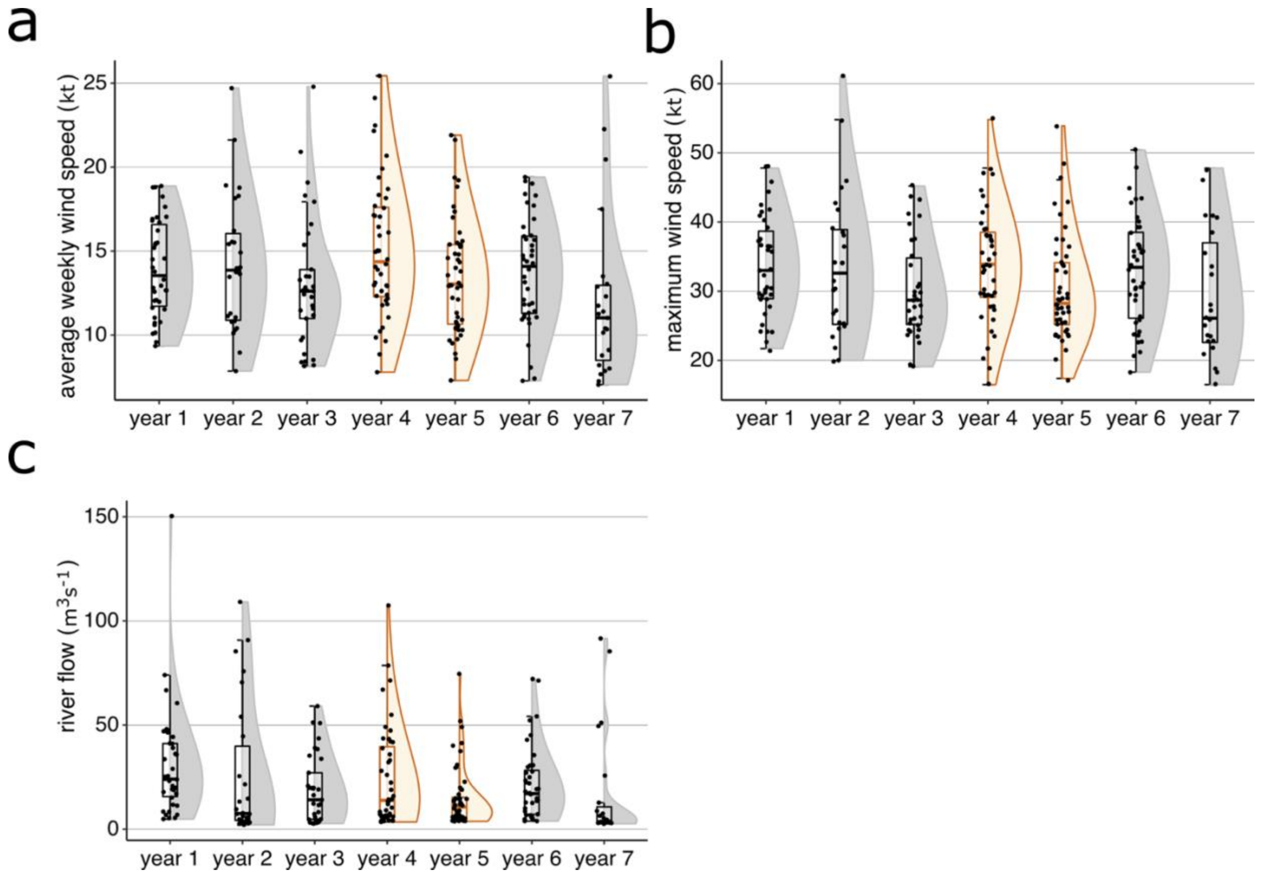


**Supplementary Figure 13. Distribution of average weekly wind speed, maximum wind speed and river flow from 2012 to 2018 at the Western English Channel. Violin plot overlapped by a boxplot and the corresponding measurements as black points.** Each group labeled as "year" in the x-axis represent a year starting and ending in the vernal equinox (March 21). Year 4 (2015-2016) and year 5 (2016-2017) are highlighted in yellow. These years correspond to the highly weekly sampling period showed in Fig S9 (between red lines in panel 'a' and panel 'b').
