## Supplementary material for "Influence of short and long term processes on SAR11 communities in open ocean and coastal systems": A list of samples is provided as Table S1

| Bermuda Atlantic Time-series | | | | | | | | | | | | |
|---|---|---|---|---|---|---|---|---|---|---|---|---|
| Date (YYYY-MM-DD) | Sample | Temperature (°C) | Salinity (p.s.u.) | O2 (μmol/kg) | NO2+NO3 (μmol/kg) | NO3 (μmol/kg) | NO2 (μmol/kg) | PO4 (μmol/kg) | SiO2 (μmol/kg) | Bacteria (cells*10^8/kg) | Turner. Chl.a (μg/L) | Reference |
| 1991-08-12 | 35-0 | 29.580 | 36.893 | 195.830 | 0 | 0 | 0 | 0 | 0.83 | 1.6 | 0.02 | Vergin et al., 2013 |
| 1991-09-12 | 36-0 | 26.420 | 36.266 | 206.390 | 0 | 0 | 0 | 0 | 0.96 | 4.01 | 0.09 | Vergin et al., 2013 |
| 1991-10-07 | 37-0 | 25.770 | 36.496 | 210.820 | 0 | 0 | 0 | 0 | 0.68 | 4.31 | - | Vergin et al., 2013 |
| 1991-11-11 | 38-0 | 23.380 | 36.598 | 217.090 | 0 | 0 | 0 | 0 | 0.69 | 4.38 | 0.02 | Vergin et al., 2013 |
| 1991-12-09 | 39-0 | 22.796 | 36.804 | 219.930 | 0 | 0 | 0 | 0 | 0.71 | 4.52 | 0.06 | Vergin et al., 2013 |
| 1992-01-09 | 40-0 | 20.742 | 36.697 | - | 0.07 | 0.03 | 0.04 | 0 | 0.7 | 4.81 | 0.27 | Vergin et al., 2013 |
| 1992-02-13 | 41-0 | 19.277 | 36.677 | 224.720 | 0.65 | 0.36 | 0.29 | 0 | 0.86 | 5.91 | 0.3 | Vergin et al., 2013 |
| 1992-03-10 | 42-0 | 19.659 | 36.659 | 229.170 | 0 | 0 | 0 | 0 | 0.68 | 7.29 | 0.11 | Vergin et al., 2013 |
| 1992-04-21 | 43-0 | 20.150 | 36.786 | 226.760 | 0 | 0 | 0 | 0 | 0.64 | 4.46 | 0.06 | Vergin et al., 2013 |
| 1992-05-18 | 44-0 | 21.867 | 36.710 | 223.750 | 0 | 0 | 0 | 0 | 0.6 | - | 0.04 | Vergin et al., 2013 |
| 1992-06-17 | 45-0 | 23.661 | 36.398 | 215.770 | 0 | 0 | 0 | 0 | 0.71 | 6.43 | 0.04 | Vergin et al., 2013 |
| 1992-07-14 | 46-0 | 26.503 | 36.370 | 206.860 | 0 | 0 | 0 | 0 | 0.76 | 4.6 | 0.03 | Vergin et al., 2013 |
| 1992-08-18 | 47-0 | 28.327 | 36.449 | 201.890 | 0 | 0 | 0 | 0.11 | 0.7 | 4.37 | 0.04 | Vergin et al., 2013 |
| 1992-09-15 | 48-0 | 26.810 | 36.299 | 202.780 | 0 | 0 | 0 | 0 | 0.76 | 7 | 0.06 | Vergin et al., 2013 |
| 1992-10-14 | 49-0 | 25.172 | 36.340 | 207.130 | 0 | 0 | 0 | 0 | 0.85 | - | 0.06 | Vergin et al., 2013 |
| 1992-11-12 | 50-0 | 23.788 | 36.540 | 211.220 | 0 | 0 | 0 | 0.04 | 0.81 | 4.93 | 0.1 | Vergin et al., 2013 |
| 1992-12-07 | 51-0 | 22.235 | 36.578 | 215.100 | 0 | 0 | 0 | 0 | 0.68 | 4.9 | 0.25 | Vergin et al., 2013 |
| 1993-01-12 | 52-0 | 21.110 | 36.660 | 222.230 | 0 | 0 | 0 | 0 | 0.72 | 4.68 | 0.12 | Vergin et al., 2013 |
| 1993-02-09 | 53-0 | 20.153 | 36.670 | 217.400 | 0 | 0 | 0 | 0 | 0.77 | - | 0.27 | Vergin et al., 2013 |
| 1993-03-10 | 54-0 | 19.279 | 36.640 | 224.190 | 0.28 | 0.19 | 0.088 | 0 | 0.87 | 4.76 | 0.27 | Vergin et al., 2013 |
| 1993-04-14 | 55-0 | 20.815 | 36.750 | - | 0 | 0 | 0 | 0 | 0.78 | - | 0.07 | Vergin et al., 2013 |
| 1993-05-09 | 56-0 | 21.550 | 36.542 | 226.410 | 0 | 0 | 0 | 0.08 | 0.71 | 5.82 | 0.1 | Vergin et al., 2013 |
| 1993-06-15 | 57-0 | 23.910 | 36.761 | 209.710 | 0 | 0 | 0 | 0 | 0.73 | 4.56 | 0.05 | Vergin et al., 2013 |
| 1993-07-13 | 58-0 | 26.790 | 36.745 | 205.700 | 0 | 0 | 0 | 0.06 | 0.78 | 5.2 | 0.04 | Vergin et al., 2013 |
| 1993-08-18 | 59-0 | 27.524 | 36.667 | 200.300 | 0 | 0 | 0 | 0.04 | 0.7 | 4.25 | 0.06 | Vergin et al., 2013 |
| 1993-10-12 | 61-0 | 26.020 | 36.435 | 204.130 | 0 | 0 | 0 | 0 | 0.83 | 4.82 | 0.08 | Vergin et al., 2013 |
| 1993-11-09 | 62-0 | 24.940 | 36.526 | 209.400 | 0 | 0 | 0 | 0 | 1.18 | 5.23 | 0.05 | Vergin et al., 2013 |
| 1993-12-07 | 63-0 | 23.300 | 36.476 | 213.500 | 0 | 0 | 0 | 0.1 | 0.82 | 3.66 | 0.04 | Vergin et al., 2013 |
| 1994-02-18 | 65-0 | 20.358 | 36.612 | 222.010 | 0 | 0 | 0 | 0 | 0.95 | 5.49 | 0.14 | Vergin et al., 2013 |
| 1997-09-12 | 108-0 | 27.037 | 36.703 | 199.560 | 0 | 0 | 0.01 | 0 | 0.94 | 3.66 | 0.03 | Vergin et al., 2013 |
| 1997-10-07 | 109-0 | 26.600 | 36.406 | 201.500 | 0 | 0 | 0 | 0 | 0.35 | 4.43 | 0.05 | Vergin et al., 2013 |
| 1997-11-12 | 110-0 | 24.158 | 36.599 | 209.100 | 0 | 0 | 0 | 0 | 0.46 | 4.84 | 0.08 | Vergin et al., 2013 |
| 1997-12-09 | 111-0 | 22.060 | 36.530 | 212.850 | 0 | 0 | 0.02 | 0 | 0.82 | 5.14 | 0.21 | Vergin et al., 2013 |
| 1998-01-13 | 112-0 | 21.890 | 36.760 | 216.140 | 0 | 0 | 0 | 0 | 0.62 | 5.27 | 0.08 | Vergin et al., 2013 |
| 1998-02-12 | 113-0 | 20.380 | 36.682 | 219.550 | 0.06 | 0.02 | 0.04 | 0 | 0.63 | 4.55 | 0.26 | Vergin et al., 2013 |
| 1998-03-12 | 114-0 | 19.590 | 36.622 | 225.700 | 0 | 0 | 0.02 | 0 | 0.65 | 3.56 | 0.27 | Vergin et al., 2013 |
| 1998-04-07 | 115-0 | 19.990 | 36.653 | 223.840 | 0 | 0 | 0.01 | 0 | 0.92 | 5.19 | 0.15 | Vergin et al., 2013 |
| 1998-05-05 | 116-0 | 20.960 | 36.668 | 222.030 | 0 | 0 | 0.01 | 0 | 0.77 | 3.99 | 0.09 | Vergin et al., 2013 |
| 1998-06-01 | 117-0 | 23.080 | 36.450 | 216.350 | 0.06 | 0.05 | 0.01 | 0 | - | 5.78 | 0.08 | Vergin et al., 2013 |
| 1998-07-07 | 118-0 | 26.730 | 36.310 | 204.060 | 0 | 0 | 0 | 0 | 0.41 | 5.29 | 0.07 | Vergin et al., 2013 |
| 1998-09-10 | 120-0 | 28.140 | 36.190 | 197.890 | 0 | 0 | 0 | 0 | 0.35 | 5.86 | 0.08 | Vergin et al., 2013 |
| 1998-12-08 | 123-0 | 23.090 | 36.560 | 213.050 | 0 | 0 | 0 | 0 | 0.55 | 4.18 | 0.07 | Vergin et al., 2013 |

| | | | | | | | | | | | | |
|---|---|---|---|---|---|---|---|---|---|---|---|---|
| 1999-01-15 | 124-0 | 21.910 | 36.500 | 217.440 | 0 | 0 | 0 | 0 | 1.03 | 5.76 | 0.08 | Vergin et al., 2013 |
| 1999-02-12 | 125-0 | 21.430 | 36.650 | 216.720 | 0 | 0 | 0 | 0 | 0.69 | 5.74 | 0.18 | Vergin et al., 2013 |
| 1999-03-23 | 126-0 | 19.480 | 36.300 | 226.300 | 0.06 | 0.06 | 0 | 0 | 0 | 2.76 | 0.41 | Vergin et al., 2013 |
| 1999-04-07 | 127-0 | 19.710 | 36.650 | 223.230 | 0.1 | 0.04 | 0.06 | 0 | 0 | - | 0.35 | Vergin et al., 2013 |
| 1999-05-04 | 128-0 | 21.483 | 36.580 | 220.500 | 0 | 0 | 0 | 0 | 0 | - | 0.07 | Vergin et al., 2013 |
| 1999-06-01 | 129-0 | 22.910 | 36.580 | 215.820 | - | - | 0 | - | 0 | 5.34 | 0.04 | Vergin et al., 2013 |
| 1999-07-06 | 130-0 | 27.150 | 36.780 | 202.150 | 0 | 0 | 0.01 | 0 | 0.72 | 3.22 | 0.04 | Vergin et al., 2013 |
| 1999-08-03 | 131-0 | 27.400 | 36.610 | 200.580 | 0 | 0 | 0 | 0 | 0.38 | 6.05 | 0.06 | Vergin et al., 2013 |
| 1999-09-15 | 132-0 | 27.790 | 36.360 | 198.370 | 0.07 | 0.07 | 0 | 0 | 0.44 | 5.46 | 0.04 | Vergin et al., 2013 |
| 1999-10-12 | 133-0 | 26.260 | 36.500 | 216.010 | 0 | 0 | 0 | 0 | 0 | 5.05 | 0.03 | Vergin et al., 2013 |
| 1999-11-09 | 134-0 | 24.490 | 36.560 | 206.900 | 0 | 0 | 0 | 0 | 0.4 | 4.81 | 0.11 | Vergin et al., 2013 |
| 1999-12-10 | 135-0 | 22.810 | 36.810 | 209.980 | 0 | 0 | 0.01 | 0 | 0.41 | 5 | 0.14 | Vergin et al., 2013 |
| 2000-01-29 | 136-0 | 20.520 | 36.750 | 215.460 | 0 | 0 | 0 | 0 | 0.46 | 4.56 | 0.22 | Vergin et al., 2013 |
| 2000-02-15 | 137-0 | 19.960 | 36.670 | 220.670 | 0 | 0 | 0 | 0 | 0.35 | 6.97 | 0.09 | Vergin et al., 2013 |
| 2000-03-14 | 138-0 | 19.360 | 36.650 | 226.050 | 0.11 | 0.11 | 0 | 0 | 0.72 | 6.51 | 0.16 | Vergin et al., 2013 |
| 2000-04-11 | 139-0 | 19.930 | 36.660 | 229.290 | 0 | 0 | 0 | 0 | 0.93 | 9.94 | 0.15 | Vergin et al., 2013 |
| 2000-05-10 | 140-0 | 22.020 | 36.600 | 222.000 | 0 | 0 | 0 | 0 | 0.81 | 6.8 | 0.05 | Vergin et al., 2013 |
| 2000-06-06 | 141-0 | 24.930 | 36.830 | 208.540 | 0 | - | - | 0 | - | 2.83 | 0.01 | Vergin et al., 2013 |
| 2000-07-11 | 142-0 | 26.510 | 36.490 | 202.980 | 0 | 0 | 0 | 0 | 0.65 | 4.19 | 0.02 | Vergin et al., 2013 |
| 2000-08-08 | 143-0 | 28.910 | 36.540 | 202.120 | 0 | 0 | 0 | 0 | 0 | 4.19 | 0.01 | Vergin et al., 2013 |
| 2000-09-13 | 144-0 | 27.840 | 36.070 | 197.570 | 0 | - | 0.01 | 0 | 0.23 | 3.97 | 0.02 | Vergin et al., 2013 |
| 2000-10-18 | 145-0 | 26.140 | 36.680 | 203.370 | - | - | - | - | 0 | 2.97 | 0.02 | Vergin et al., 2013 |
| 2000-11-14 | 146-0 | 23.360 | 36.660 | 210.340 | 0 | 0 | 0 | 0 | 0.34 | 3.1 | 0.11 | Vergin et al., 2013 |
| 2000-12-12 | 147-0 | 21.990 | 36.600 | 214.810 | 0.06 | 0.06 | 0 | - | 0.41 | 3.64 | 0.02 | Vergin et al., 2013 |
| 2001-01-30 | 148-0 | 20.140 | 36.610 | 219.570 | 0.06 | 0.03 | 0.03 | 0 | 0.43 | 3.43 | 0.27 | Vergin et al., 2013 |
| 2001-02-20 | 149-0 | 19.930 | 36.650 | 222.190 | 0 | 0 | 0.01 | 0 | 0.24 | 4.7 | 0.24 | Vergin et al., 2013 |
| 2001-03-09 | 150-0 | 19.680 | 36.690 | 220.690 | 0.06 | 0.03 | 0.03 | - | - | 3.56 | 0.21 | Vergin et al., 2013 |
| 2001-04-04 | 151-0 | 20.130 | 36.720 | 224.570 | 0 | 0 | 0.01 | 0 | 1.25 | 4.84 | 0.05 | Vergin et al., 2013 |
| 2001-05-15 | 152-0 | 21.340 | 36.780 | 219.520 | 0.23 | 0.22 | 0.01 | 0 | 0.47 | 3.66 | 0.04 | Vergin et al., 2013 |
| 2001-06-05 | 153-0 | 24.470 | 36.850 | 210.870 | 0 | 0 | 0 | 0 | 1.32 | 3.05 | 0.05 | Vergin et al., 2013 |
| 2001-06-01 | 153-0 | 24.470 | 36.850 | 210.870 | 0 | 0 | 0 | 0 | 1.32 | 3.05 | 0.05 | Vergin et al., 2013 |
| 2001-07-18 | 154-0 | 26.500 | 36.500 | 208.420 | 0 | 0 | 0 | 0 | 1.14 | 4.04 | 0.07 | Vergin et al., 2013 |
| 2001-08-07 | 155-0 | 27.550 | 36.530 | 201.250 | 0 | 0 | 0 | 0 | 1.06 | 3.56 | 0.06 | Vergin et al., 2013 |
| 2001-09-12 | 156-0 | 27.560 | 36.520 | 202.340 | 0 | 0 | 0 | 0 | 1.22 | 3.04 | 0.06 | Vergin et al., 2013 |
| 2001-10-16 | 157-0 | 24.680 | 36.440 | 209.770 | 0 | 0 | 0 | 0 | 0.53 | 4.63 | 0.1 | Vergin et al., 2013 |
| 2001-11-09 | 158-0 | 24.370 | 36.550 | 209.940 | 0 | 0 | 0 | 0 | 0.46 | 3.84 | 0.06 | Vergin et al., 2013 |
| 2001-12-11 | 159-0 | 22.280 | 36.620 | 215.590 | 0 | 0 | 0 | 0.04 | 0.66 | 3.7 | 0.12 | Vergin et al., 2013 |
| 2002-01-01 | 160-0 | - | - | - | - | - | - | - | - | - | - | Vergin et al., 2013 |
| 2002-02-13 | 161-0 | 20.350 | 36.690 | 221.270 | 0 | 0 | 0.02 | 0.04 | 0.47 | 3.2 | 0.26 | Vergin et al., 2013 |
| 2002-03-19 | 162-0 | 19.940 | 36.670 | 226.390 | 0 | 0 | 0.01 | 0 | 0.66 | 4.72 | 0.13 | Vergin et al., 2013 |
| 2002-04-16 | 163-0 | 21.250 | 36.760 | 221.310 | 0 | 0 | 0 | 0.04 | 0.7 | 3.62 | 0.05 | Vergin et al., 2013 |
| 2002-05-14 | 164-0 | 21.720 | 36.520 | 222.860 | 0 | 0 | 0 | 0 | - | 3.68 | 0.07 | Vergin et al., 2013 |
| 2002-06-04 | 165-0 | 23.490 | 36.470 | 215.140 | 0 | 0 | 0 | 0 | 0.25 | 3.71 | 0.05 | Vergin et al., 2013 |
| 2002-07-02 | 166-0 | 26.870 | 36.620 | 202.570 | 0 | 0 | 0 | 0 | 0.81 | 1.97 | 0.05 | Vergin et al., 2013 |
| 2002-08-20 | 167-0 | 27.870 | 36.320 | 200.520 | 0 | 0 | 0 | 0 | 1.32 | 4.64 | - | Vergin et al., 2013 |

| | | | | | | | | | | | | |
|---|---|---|---|---|---|---|---|---|---|---|---|---|
| 2002-09-17 | 168-0 | 26.990 | 36.210 | 198.250 | 0 | 0 | 0 | 0 | 0 | 4.48 | 0.06 | Vergin et al., 2013 |
| 2002-10-08 | 169-0 | 26.410 | 36.190 | 202.930 | 0 | 0 | 0 | 0.06 | 0.27 | 4.49 | 0.02 | Vergin et al., 2013 |
| 2002-11-13 | 170-0 | 24.680 | 36.530 | 209.910 | 0 | 0 | 0.01 | 0 | 0.92 | 4.16 | 0.07 | Vergin et al., 2013 |
| 2002-12-10 | 171-0 | 23.400 | 36.660 | 211.680 | 0 | 0 | 0 | 0 | 0.7 | 3.98 | 0.17 | Vergin et al., 2013 |
| 2003-01-22 | 172-0 | 20.820 | 36.720 | 218.410 | 0 | 0 | 0.03 | 0 | 0.63 | 3.9 | 0.14 | Vergin et al., 2013 |
| 2003-02-21 | 173-0 | 20.600 | 36.740 | 222.010 | 0 | 0 | 0 | 0 | 1.08 | 4.27 | 0.16 | Vergin et al., 2013 |
| 2003-03-21 | 174-0 | 19.990 | 36.670 | 224.820 | 0 | 0 | 0.01 | 0 | 1.44 | 4.86 | 0.06 | Vergin et al., 2013 |
| 2003-04-22 | 175-0 | 21.770 | 36.580 | 219.310 | 0 | 0 | 0 | 0 | 0.57 | 4.78 | 0.06 | Vergin et al., 2013 |
| 2003-05-20 | 176-0 | 22.700 | 36.680 | 216.600 | 0 | 0 | 0 | 0 | 0.63 | 4.85 | 0.06 | Vergin et al., 2013 |
| 2003-06-01 | 177-0 | 26.760 | 36.390 | 207.530 | 0 | 0 | 0 | 0 | 0.6 | 3.98 | 0.02 | Vergin et al., 2013 |
| 2003-07-15 | 178-0 | 27.830 | 36.450 | 204.520 | - | - | - | - | - | 4.44 | 0.03 | Vergin et al., 2013 |
| 2003-08-12 | 179-0 | 28.860 | 36.640 | 198.980 | 0 | 0 | 0 | 0 | 0.7 | 4.48 | 0.05 | Vergin et al., 2013 |
| 2003-09-19 | 180-0 | 26.170 | 36.470 | 210.010 | 0 | 0 | 0 | 0 | 0.91 | 4.77 | 0.09 | Vergin et al., 2013 |
| 2003-10-07 | 181-0 | 26.350 | 36.392 | 201.800 | 0 | 0 | 0 | 0.04 | 0.94 | 3.93 | 0.08 | Vergin et al., 2013 |
| 2003-12-02 | 183-0 | 24.210 | 36.800 | 208.850 | 0 | 0 | 0 | 0 | 0.99 | 4.3 | 0.14 | Vergin et al., 2013 |
| 2004-01-27 | 184-0 | 20.300 | 36.693 | 219.800 | 0.11 | 0.11 | 0 | 0 | 0.9 | 4.38 | 0.39 | Vergin et al., 2013 |
| 2016-03-08 | 10321-1_S39 | 21.151 | 36.671 | 314.200 | 0 | 0 | 0.02 | 0 | 0 | 7.7 | 0.11 | BATS this study |
| 2016-03-25 | 10322-1_S51 | 20.899 | 36.624 | - | - | - | - | - | - | - | 0.1 | BATS this study |
| 2016-04-04 | 20322-1_S63 | 21.533 | 36.664 | 218.200 | - | - | - | - | - | - | 0.08 | BATS this study |
| 2016-04-15 | SC1_S91 | 21.350 | 36.701 | 221.100 | 0 | 0 | 0 | 0 | 0.35 | 6.8 | 0.06 | BATS this study |
| 2016-05-03 | SC23_S113 | 21.789 | 36.647 | 220.900 | 0 | 0 | 0.01 | 0.05 | 0 | 6.4 | 0.08 | BATS this study |
| 2016-07-10 | C7_N1_S112 | 26.674 | 36.336 | 204.786 | 0.04 | - | - | 0.02 | - | 7 | - | BATS this study |
| 2016-08-17 | 10327-1_S89 | 29.384 | 36.508 | 196.600 | 0 | - | - | 0 | 0.6 | 3.3 | 0.03 | BATS this study |
| 2016-09-22 | 10328-1_S101 | 28.776 | 36.706 | 195.400 | 0 | 0 | 0 | 0 | 0.68 | 3 | 0.04 | BATS this study |
| 2016-10-20 | BS-1_S62 | 26.158 | 36.815 | 206.000 | 0 | 0 | 0.02 | 0 | 0 | 4.2 | 0.07 | BATS this study |
| 2016-11-21 | BS-13_S74 | 22.952 | 36.629 | - | 0 | - | - | 0 | 0 | 3.8 | 0.1 | BATS this study |
| 2016-12-15 | BS-25_S86 | 21.765 | 36.557 | 216.700 | 0 | 0 | 0.02 | 0 | 0 | 4.5 | 0.11 | BATS this study |
| 2017-01-07 | BS-37_S98 | 21.654 | 36.628 | 219.366 | 0 | 0 | 0.01 | 0 | 0.91 | 6.62 | 0.11 | BATS this study |
| 2017-02-19 | BS-49_S110 | 21.107 | 36.777 | 218.267 | - | - | - | - | - | 5.15 | 0.16 | BATS this study |
| 2017-04-10 | BS-61_S122 | 20.454 | 36.740 | 223.870 | 0.19 | 0 | 0.19 | 0.07 | 0.97 | 4.38 | 0.17 | BATS this study |
| 2017-04-18 | BS-73_S134 | 21.277 | 36.845 | 220.859 | - | - | - | - | - | 4.23 | 0.05 | BATS this study |
| 2017-05-10 | BS-81_S142 | 23.187 | 36.834 | 214.562 | 0 | 0 | 0 | 0 | 0.51 | 4.34 | 0.03 | BATS this study |
| 2017-05-30 | BS93_S1 | 24.090 | 36.742 | 215.308 | - | - | - | - | - | 4.14 | 0.05 | BATS this study |
| 2017-06-14 | BS101_S9 | 25.936 | 36.738 | 209.371 | - | - | - | - | - | 6.87 | 0.05 | BATS this study |
| 2017-07-19 | BS113_S21 | 28.230 | 36.612 | 200.337 | 0 | 0 | 0 | 0 | 0.39 | 6.2 | 0.05 | BATS this study |
| 2017-08-17 | BS124_S32 | 29.054 | 36.832 | 196.921 | 0 | 0 | 0.22 | 0.04 | 0.85 | 6.01 | 0.04 | BATS this study |
| 2017-09-14 | BS135_S43 | 28.391 | 36.522 | 196.308 | 0 | 0 | 0.01 | 0 | 0.97 | 6.48 | 0.05 | BATS this study |
| 2017-10-17 | BS147_S55 | 25.944 | 36.518 | 205.366 | 0 | 0 | 0.01 | 0 | 1.03 | 6.3 | 0.06 | BATS this study |
| 2017-11-18 | BS159_S67 | 24.471 | 36.538 | 206.241 | 0 | 0 | 0.01 | 0 | 1.1 | 4.63 | 0.05 | BATS this study |
| 2017-12-14 | BS171_S79 | 23.132 | 36.664 | 210.645 | 0 | 0 | 0.01 | 0 | 0.86 | 4.14 | 0.12 | BATS this study |
| 2018-01-17 | BS_183_S21 | 21.787 | 36.681 | 216.803 | - | - | - | 0 | 0.88 | 4.44 | 0.07 | BATS this study |
| 2018-02-12 | BS_195_S33 | 21.065 | 36.793 | 221.450 | 0 | 0 | 0 | 0 | 0.79 | 6.35 | 0.09 | BATS this study |
| 2018-02-26 | BS_207_S45 | 21.333 | 36.805 | 221.305 | 0 | 0 | 0.01 | 0 | 0.81 | - | 0.02 | BATS this study |
| 2018-03-25 | BS_215_S53 | 20.186 | 36.781 | 221.793 | - | - | - | - | - | 5.3 | 0.16 | BATS this study |
| 2018-04-16 | BS_227_S65 | 20.880 | 36.782 | 225.881 | - | - | - | - | - | 1.85 | 0.05 | BATS this study |

| | | | | | | | | | | | | |
|---|---|---|---|---|---|---|---|---|---|---|---|---|
| 2018-04-25 | BS_239_S77 | 20.993 | 36.796 | 228.764 | 0 | 0 | 0.01 | 0 | 0.83 | 6.75 | 0.09 | BATS this study |
| 2018-05-25 | BS_247_S85 | 24.414 | 36.690 | 213.043 | 0 | 0 | 0 | 0 | 0.79 | 5.17 | 0.03 | BATS this study |
| 2018-06-14 | BS_259_S97 | 25.415 | 36.550 | 207.924 | 0 | 0 | 0 | 0 | 0.9 | 3.81 | 0.04 | BATS this study |
| 2018-07-21 | BS271_S109 | 26.992 | 36.534 | 201.715 | 0 | 0 | 0 | 0 | 0.87 | 3.89 | 0.02 | BATS this study |
| 2018-08-14 | BS283_S121 | 28.358 | 36.591 | 197.078 | 0 | 0 | 0 | 0 | 1.01 | 3.77 | 0.01 | BATS this study |
| 2018-09-12 | BS295_S133 | 28.721 | 36.665 | 200.157 | 0 | 0 | 0 | 0 | 0.88 | 4.09 | 0.03 | BATS this study |
| 2018-10-21 | BS307_S145 | 25.713 | 36.960 | 204.511 | 0 | 0 | 0 | 0 | 0.81 | 4.77 | 0.04 | BATS this study |
| 2018-11-07 | BS319_S157 | 24.514 | 36.608 | 209.879 | 0 | 0 | 0 | 0 | 0.94 | 6.8 | 0.06 | BATS this study |
| 2018-12-15 | BS330_S167 | 22.336 | 36.793 | 215.435 | 0 | 0 | 0 | 0 | 0.81 | - | 0.07 | BATS this study |

**Wester English Channel Time-series**

| Date (YYYY-MM-DD) | Sample | Temperature (°C) | Salinity (p.s.u.) | O2 (µmol/kg) | NO2+NO3 (µmol/kg) | NO2 (µmol/kg) | NH4 (µmol/kg) | PO4 (µmol/kg) | SiO2 (µmol/kg) | Bacteria (cells*10^8/kg) | Chl.a (µg/L) | Reference |
|---|---|---|---|---|---|---|---|---|---|---|---|---|
| 2012-02-06 | KT16S193 | 9.486 | 35.290 | 357.648 | 6.910 | 0.255 | - | 0.440 | 3.325 | - | 0.310 | WEC this study |
| 2012-02-13 | KT16S194 | 9.210 | 35.110 | 272.024 | 7.610 | 0.305 | 0.205 | 0.455 | 3.910 | - | 0.400 | WEC this study |
| 2012-02-20 | KT16S195 | 9.129 | 35.166 | 273.651 | 7.603 | 0.310 | 0.130 | 0.540 | 3.868 | - | 0.610 | WEC this study |
| 2012-02-27 | KT16S196 | 9.366 | 35.117 | 275.953 | 8.100 | 0.373 | - | 0.525 | 3.563 | - | 0.680 | WEC this study |
| 2012-03-12 | KT16S197 | 9.571 | 35.255 | 274.137 | 6.103 | 0.395 | 0.373 | 0.450 | 3.300 | - | 0.860 | WEC this study |
| 2012-03-19 | KT16S198 | 9.559 | 35.162 | 282.236 | 5.795 | 0.330 | 0.168 | 0.378 | 3.133 | - | 1.770 | WEC this study |
| 2012-03-26 | KT16S199 | 9.762 | 35.302 | 278.651 | 5.345 | 0.278 | 0.498 | 0.350 | - | - | 1.770 | WEC this study |
| 2012-04-02 | KT16S200 | 10.102 | 35.327 | 294.946 | 2.695 | 0.125 | 0.270 | 0.240 | 1.175 | - | 4.350 | WEC this study |
| 2012-04-16 | KT16S201 | 10.144 | 35.332 | 301.453 | 0.295 | 0.058 | 0.230 | 0.125 | 0.215 | - | 3.610 | WEC this study |
| 2012-04-23 | KT16S202 | 10.337 | 35.257 | 281.601 | 0.105 | 0.060 | 0.200 | 0.085 | 0.268 | - | 2.730 | WEC this study |
| 2012-04-30 | KT16S203 | 10.314 | 35.182 | 273.332 | 0.320 | 0.065 | 0.245 | 0.133 | 1.393 | - | 2.120 | WEC this study |
| 2012-05-07 | KT16S204 | 10.902 | 35.259 | 272.072 | 0.413 | 0.050 | 0.798 | 0.113 | 1.583 | - | 1.130 | WEC this study |
| 2012-05-21 | KT16S205 | 11.390 | 35.124 | 317.496 | 0.078 | 0.013 | - | 0.040 | 0.325 | - | 2.450 | WEC this study |
| 2012-05-28 | KT16S206 | - | - | - | - | - | - | - | - | - | - | WEC this study |
| 2012-06-11 | KT16S207 | 12.794 | 35.076 | 272.388 | 0.095 | 0.020 | 0.395 | 0.075 | 0.348 | - | 3.130 | WEC this study |
| 2012-06-25 | KT16S208 | 13.941 | 34.638 | 264.453 | - | - | - | - | - | - | 2.450 | WEC this study |
| 2012-07-02 | KT16S209 | 14.074 | 35.085 | 251.719 | 0.085 | 0.010 | 0.168 | 0.030 | 0.185 | - | 1.390 | WEC this study |
| 2012-07-09 | KT16S210 | 15.080 | 34.534 | 253.191 | 2.670 | 0.070 | 0.338 | 0.080 | 1.360 | - | 1.480 | WEC this study |
| 2012-07-23 | KT16S211 | 15.973 | 35.069 | 263.852 | 0.040 | 0.000 | 0.000 | 0.023 | 0.105 | - | 0.630 | WEC this study |
| 2012-07-30 | KT16S212 | 15.053 | 35.089 | 267.397 | 0.000 | 0.000 | 0.635 | 0.038 | 0.525 | - | 2.050 | WEC this study |
| 2012-08-06 | KT16S213 | 15.419 | 35.171 | 235.778 | 0.115 | 0.000 | 0.615 | 0.060 | 0.395 | - | 1.610 | WEC this study |
| 2012-08-13 | KT16S214 | 17.382 | 35.201 | 243.745 | 0.070 | 0.015 | 0.678 | 0.045 | 0.368 | - | 0.570 | WEC this study |
| 2012-08-20 | KT16S215 | 17.806 | 34.961 | 231.600 | 0.950 | 0.095 | 1.210 | 0.160 | 1.973 | - | 1.100 | WEC this study |
| 2012-09-17 | KT16S216 | 16.117 | 35.150 | 247.453 | 0.055 | 0.020 | 0.655 | 0.083 | 0.303 | - | 1.340 | WEC this study |
| 2012-10-08 | KT16S217 | 14.783 | 35.133 | 241.796 | 1.900 | 0.523 | 0.310 | 0.280 | 1.950 | - | 2.000 | WEC this study |
| 2012-10-15 | KT16S218 | - | - | - | 5.835 | 1.110 | 0.175 | 0.355 | 3.580 | - | 1.080 | WEC this study |
| 2012-10-29 | KT16S219 | 14.035 | 35.083 | 235.807 | 4.153 | 1.025 | 0.645 | 0.335 | 3.760 | - | 0.580 | WEC this study |
| 2012-11-12 | KT16S220 | 13.002 | 35.046 | 243.352 | 4.708 | 0.513 | 0.090 | 0.370 | 4.038 | - | 0.660 | WEC this study |
| 2012-11-26 | KT16S221 | 11.788 | 34.257 | 254.687 | 8.885 | 0.210 | 0.210 | 0.478 | 5.915 | - | 0.470 | WEC this study |
| 2012-12-03 | KT16S222 | - | - | - | 6.640 | 0.145 | 0.080 | 0.440 | 4.775 | - | 0.600 | WEC this study |
| 2012-12-10 | KT16S223 | 11.460 | 34.766 | 256.580 | 6.585 | 0.155 | 0.340 | 0.430 | 5.510 | - | 0.620 | WEC this study |
| 2012-12-17 | KT16S224 | - | - | - | 7.048 | 0.170 | - | 0.470 | 5.485 | - | 0.550 | WEC this study |
| 2013-01-07 | KT16S225 | 10.625 | 34.507 | 272.520 | 8.435 | 0.095 | - | 0.535 | 5.265 | - | 0.650 | WEC this study |

| | | | | | | | | | | | | |
|---|---|---|---|---|---|---|---|---|---|---|---|---|
| 2013-01-11 | KT16S255 | - | - | - | - | - | - | - | - | - | - | WEC this study |
| 2013-01-21 | KT16S226 | - | - | - | 7.015 | 0.070 | 0.465 | 0.498 | 4.940 | - | 0.330 | WEC this study |
| 2013-02-04 | KT16S227 | - | - | - | - | - | - | - | - | - | 0.360 | WEC this study |
| 2013-02-11 | KT16S228 | 9.099 | 34.662 | 273.129 | 9.528 | 0.210 | 0.080 | 0.520 | 5.778 | - | 0.500 | WEC this study |
| 2013-02-18 | KT16S229 | - | - | - | 7.928 | 0.180 | 0.140 | 0.565 | 4.913 | - | 0.410 | WEC this study |
| 2013-02-25 | KT16S230 | 8.352 | 34.830 | 281.505 | 8.198 | 0.190 | 0.100 | 0.510 | 5.843 | - | 0.410 | WEC this study |
| 2013-03-11 | KT16S231 | 8.217 | 35.156 | 283.282 | 6.635 | 0.180 | 0.280 | 0.425 | 4.100 | - | 0.660 | WEC this study |
| 2013-03-18 | KT16S232 | 7.985 | 34.760 | 289.499 | 7.915 | 0.190 | 0.338 | 0.500 | 5.123 | - | 1.090 | WEC this study |
| 2013-04-01 | KT16S233 | 7.629 | 35.106 | 286.052 | 6.988 | 0.230 | - | 0.480 | 3.845 | - | 0.520 | WEC this study |
| 2013-04-29 | KT16S234 | 8.220 | 35.035 | 289.391 | 6.845 | 0.190 | 0.530 | 0.430 | 3.558 | - | 3.340 | WEC this study |
| 2013-05-06 | KT16S235 | - | - | - | 2.967 | 0.135 | 0.363 | 0.180 | 2.063 | - | 3.150 | WEC this study |
| 2013-05-13 | KT16S236 | - | - | - | 2.023 | 0.090 | 0.713 | 0.150 | 0.363 | - | 2.070 | WEC this study |
| 2013-05-20 | KT16S237 | 10.476 | 34.920 | 317.571 | 0.220 | 0.030 | 0.293 | 0.063 | 0.298 | - | 2.650 | WEC this study |
| 2013-05-27 | KT16S238 | 10.365 | 35.191 | 316.298 | 0.055 | 0.000 | 0.333 | 0.055 | 0.670 | - | 0.280 | WEC this study |
| 2013-06-03 | KT16S239 | 11.573 | 35.169 | 313.445 | 0.015 | 0.000 | 0.300 | 0.120 | 0.850 | - | 0.100 | WEC this study |
| 2013-06-10 | KT16S240 | - | - | - | - | - | - | - | - | - | 0.510 | WEC this study |
| 2013-06-17 | KT16S241 | - | - | - | 0.060 | 0.010 | 0.123 | 0.055 | 0.800 | - | 0.740 | WEC this study |
| 2013-06-24 | KT16S242 | 12.311 | 35.029 | 272.140 | 0.000 | 0.000 | 0.258 | 0.045 | - | - | 1.260 | WEC this study |
| 2013-07-01 | KT16S243 | 13.139 | 35.165 | 271.387 | 0.073 | 0.025 | 0.047 | 0.040 | 1.330 | - | 0.460 | WEC this study |
| 2013-07-08 | KT16S244 | 15.741 | 35.166 | 279.836 | 0.027 | 0.023 | 0.020 | 0.010 | 0.203 | - | 0.440 | WEC this study |
| 2013-07-15 | KT16S245 | 17.240 | 35.249 | 270.267 | 0.037 | 0.010 | 0.000 | 0.010 | 0.188 | - | 0.790 | WEC this study |
| 2013-09-02 | KT16S246 | 16.392 | 35.130 | 197.876 | 0.013 | 0.000 | - | 0.020 | 1.420 | - | 0.960 | WEC this study |
| 2013-09-23 | KT16S247 | 15.134 | 35.164 | 242.623 | 1.813 | 0.500 | 0.230 | 0.163 | 1.803 | - | - | WEC this study |
| 2013-09-30 | KT16S248 | - | - | - | 2.053 | 0.930 | 0.350 | 0.210 | 2.473 | - | 0.980 | WEC this study |
| 2013-10-07 | KT16S249 | 16.226 | 35.132 | 251.965 | 1.198 | 0.555 | 0.130 | 0.195 | 3.150 | - | 1.750 | WEC this study |
| 2013-11-04 | KT16S250 | 13.930 | 34.968 | 233.630 | 6.413 | 0.605 | 0.130 | 0.393 | 3.918 | - | 0.500 | WEC this study |
| 2013-11-11 | KT16S251 | - | - | - | 6.640 | 0.505 | 0.105 | 0.348 | 3.925 | - | 0.780 | WEC this study |
| 2013-11-18 | KT16S252 | 13.103 | 34.783 | 245.186 | 7.233 | 0.285 | 0.180 | 0.385 | 4.335 | - | 0.510 | WEC this study |
| 2013-12-09 | KT16S253 | - | - | - | 6.853 | 0.270 | 0.098 | 0.415 | 3.150 | - | 0.330 | WEC this study |
| 2013-12-16 | KT16S254 | 11.636 | 35.066 | 259.403 | 7.028 | 0.150 | 0.233 | 0.438 | 4.288 | - | 0.510 | WEC this study |
| 2014-01-20 | KT16S256 | 9.531 | 33.459 | 269.022 | 16.230 | 0.260 | 1.033 | 0.710 | 7.953 | - | 0.320 | WEC this study |
| 2014-01-27 | KT16S257 | 9.802 | 34.840 | 265.693 | 9.983 | 0.190 | 0.570 | 0.590 | 5.253 | - | 0.310 | WEC this study |
| 2014-02-10 | KT16S258 | - | - | - | - | - | - | - | - | - | - | WEC this study |
| 2014-03-03 | KT16S259 | 8.895 | 34.869 | 279.671 | 9.850 | 0.130 | 0.505 | 0.658 | 5.728 | - | 0.350 | WEC this study |
| 2014-03-24 | KT16S260 | 9.397 | 35.067 | 277.514 | 8.730 | 0.230 | 0.423 | 0.625 | 5.350 | - | 0.410 | WEC this study |
| 2014-03-31 | KT16S261 | 9.484 | 35.093 | 275.929 | 8.990 | 0.320 | 0.430 | 0.610 | 4.958 | - | 0.410 | WEC this study |
| 2014-04-07 | KT16S262 | 10.209 | 34.842 | 246.134 | 9.020 | 0.340 | 0.373 | 0.495 | 4.540 | - | 0.500 | WEC this study |
| 2014-04-14 | KT16S263 | 10.423 | 34.625 | 308.103 | 4.403 | 0.310 | 0.313 | 0.205 | 0.988 | - | 3.840 | WEC this study |
| 2014-05-12 | KT16S264 | 10.868 | 35.277 | 275.149 | 3.285 | 0.135 | 1.085 | 0.395 | 1.605 | - | 1.870 | WEC this study |
| 2014-05-19 | KT16S265 | 12.405 | 35.151 | 224.350 | 0.015 | 0.000 | 0.400 | 0.100 | 0.463 | - | 0.420 | WEC this study |
| 2014-06-09 | KT16S266 | 14.068 | 35.178 | - | 0.020 | 0.008 | 0.533 | 0.040 | 0.080 | - | 0.410 | WEC this study |
| 2014-06-16 | KT16S267 | 15.620 | 35.214 | - | 0.025 | 0.000 | 0.315 | 0.028 | 0.095 | - | 0.490 | WEC this study |
| 2014-06-23 | KT16S268 | 17.200 | 35.234 | - | 0.095 | 0.010 | 0.177 | 0.105 | 0.143 | - | 0.370 | WEC this study |
| 2014-06-30 | KT16S269 | 16.093 | 35.099 | 254.968 | 0.025 | 0.005 | 0.198 | 0.018 | 0.343 | - | 0.920 | WEC this study |
| 2014-07-07 | KT16S270 | 17.071 | 35.097 | 249.579 | 0.035 | 0.015 | 0.168 | 0.020 | 0.060 | - | 0.630 | WEC this study |

| | | | | | | | | | | | | |
|---|---|---|---|---|---|---|---|---|---|---|---|---|
| 2014-07-14 | KT16S271 | 15.938 | 35.090 | 245.958 | 0.000 | 0.000 | 0.490 | 0.060 | 0.703 | - | 0.520 | WEC this study |
| 2014-07-21 | KT16S272 | 17.109 | 35.095 | 270.438 | 0.015 | 0.010 | 0.253 | 0.040 | 0.188 | - | 0.450 | WEC this study |
| 2014-08-18 | KT16S273 | 15.188 | 35.079 | 246.912 | 1.145 | 0.255 | 0.265 | 0.125 | 2.205 | - | 1.020 | WEC this study |
| 2014-08-25 | KT16S274 | 15.689 | 35.093 | 245.194 | 0.400 | 0.085 | 0.380 | 0.055 | 1.803 | - | 0.580 | WEC this study |
| 2014-09-08 | KT16S275 | 16.762 | 34.545 | 258.082 | 2.020 | 0.015 | 0.020 | 0.015 | - | - | 0.350 | WEC this study |
| 2014-09-15 | KT16S276 | - | - | - | 1.430 | 0.550 | 0.433 | 0.135 | - | - | 0.490 | WEC this study |
| 2014-09-22 | KT16S277 | 17.024 | 35.133 | 251.638 | 0.158 | 0.035 | 0.150 | 0.055 | 1.883 | - | 0.410 | WEC this study |
| 2014-09-29 | KT16S278 | 17.242 | 35.160 | 238.548 | 0.580 | 0.140 | 0.273 | 0.118 | 2.335 | - | 0.510 | WEC this study |
| 2014-10-13 | KT16S279 | 16.077 | 35.163 | 225.881 | 2.518 | 0.980 | 0.545 | 0.280 | 3.145 | - | 0.290 | WEC this study |
| 2014-10-20 | KT16S280 | - | - | - | - | - | - | - | - | - | 0.240 | WEC this study |
| 2014-10-24 | KT16S281 | - | - | - | - | - | - | - | - | - | - | WEC this study |
| 2014-11-03 | KT16S282 | 14.469 | 34.896 | 235.777 | 6.278 | 0.730 | 0.253 | 0.428 | 4.717 | - | 0.350 | WEC this study |
| 2014-11-17 | KT16S283 | 13.392 | 34.059 | 244.412 | 9.133 | 0.475 | 0.345 | 0.435 | 5.580 | - | 0.230 | WEC this study |
| 2014-11-24 | KT16S284 | 13.600 | 35.223 | 240.879 | 4.850 | 0.317 | 0.143 | 0.373 | 2.763 | - | 0.230 | WEC this study |
| 2014-12-01 | KT16S285 | 13.186 | 35.158 | 242.025 | 5.490 | 0.143 | 0.093 | 0.410 | 2.940 | - | 0.150 | WEC this study |
| 2014-12-08 | KT16S286 | 12.478 | 35.120 | 247.546 | 6.143 | 0.190 | 0.085 | 0.398 | 3.100 | - | 0.170 | WEC this study |
| 2014-12-15 | KT16S287 | - | - | - | 6.063 | 0.105 | 0.128 | 0.418 | 3.497 | - | 0.110 | WEC this study |
| 2015-01-05 | KT16S288 | 11.233 | 35.143 | 260.775 | 6.965 | 0.095 | 0.100 | 0.470 | 4.153 | - | 0.130 | WEC this study |
| 2015-01-19 | KT16S289 | 10.972 | 35.262 | 259.138 | 6.795 | 0.130 | 0.140 | 0.465 | 3.100 | - | 0.170 | WEC this study |
| 2015-02-02 | KT16S290 | 10.015 | 35.236 | 262.651 | 7.278 | 0.165 | 0.103 | 0.445 | 3.390 | - | 0.200 | WEC this study |
| 2015-02-09 | KT16S291 | 9.339 | 35.194 | 267.644 | 7.403 | 0.250 | 0.225 | 0.440 | 3.420 | - | 0.270 | WEC this study |
| 2015-02-16 | KT16S292 | 9.237 | 35.208 | 270.166 | 7.515 | 0.190 | 0.238 | 0.460 | 3.420 | - | 0.250 | WEC this study |
| 2015-03-09 | KT16S293 | 9.225 | 35.146 | 285.271 | 5.143 | 0.198 | - | 0.323 | 1.915 | - | 1.650 | WEC this study |
| 2015-03-16 | KT16S294 | - | - | - | 4.810 | 0.180 | 0.343 | 0.333 | 1.358 | - | 2.030 | WEC this study |
| 2015-03-23 | KT16S295 | 9.137 | 35.249 | 284.623 | 3.868 | 0.140 | 0.285 | 0.283 | 0.908 | - | 1.440 | WEC this study |
| 2015-04-06 | KT16S296 | 9.487 | 35.239 | 282.466 | 2.910 | 0.120 | 0.710 | 0.248 | 1.178 | - | 1.790 | WEC this study |
| 2015-04-20 | KT16S297 | 10.182 | 35.219 | 282.821 | 1.318 | 0.070 | 0.905 | 0.158 | 1.510 | - | 0.740 | WEC this study |
| 2015-04-27 | KT16S298 | 10.411 | 35.298 | 290.690 | 0.015 | 0.000 | 0.173 | 0.060 | 1.013 | - | 1.180 | WEC this study |
| 2015-05-04 | KT16S299 | 10.877 | 35.189 | 268.598 | 0.275 | 0.015 | - | 0.095 | 1.213 | - | 2.040 | WEC this study |
| 2015-05-11 | KT16S300 | 11.657 | 35.092 | 268.719 | 0.615 | 0.035 | - | 0.130 | 1.480 | - | 2.260 | WEC this study |
| 2015-05-18 | KT16S301 | 11.214 | 35.232 | 268.879 | 0.160 | 0.015 | - | 0.105 | 0.730 | - | 3.140 | WEC this study |
| 2015-05-27 | KT16S302 | - | - | - | - | - | - | - | - | - | - | WEC this study |
| 2015-06-01 | KT16S303 | 11.497 | 35.305 | 260.190 | 0.070 | 0.005 | - | 0.120 | 0.835 | - | 2.420 | WEC this study |
| 2015-06-08 | KT16S304 | 12.600 | 35.202 | 272.638 | 0.000 | 0.000 | - | 0.075 | 0.330 | - | 1.390 | WEC this study |
| 2015-06-15 | KT16S305 | 13.306 | 35.283 | 267.371 | 0.000 | 0.000 | 0.193 | 0.035 | 0.295 | - | 0.590 | WEC this study |
| 2015-06-22 | KT16S306 | 13.634 | 35.266 | 267.581 | 0.000 | 0.000 | 0.218 | 0.065 | 0.225 | - | 0.900 | WEC this study |
| 2015-06-29 | KT16S307 | 15.582 | 35.283 | 258.867 | 0.010 | 0.005 | - | 0.045 | - | - | 0.430 | WEC this study |
| 2015-07-06 | KT16S308 | 15.728 | 35.296 | 257.009 | 0.010 | 0.010 | - | 0.030 | 0.165 | - | 0.860 | WEC this study |
| 2015-07-13 | KT16S309 | 15.170 | 35.236 | 258.235 | 0.007 | 0.000 | 0.070 | 0.030 | 0.558 | - | 1.520 | WEC this study |
| 2015-07-20 | KT16S310 | 16.173 | 35.250 | 269.224 | 0.000 | 0.000 | 0.000 | 0.000 | 0.625 | - | 0.910 | WEC this study |
| 2015-07-28 | KT16S311 | - | - | - | - | - | - | - | - | - | - | WEC this study |
| 2015-08-03 | KT16S312 | 15.664 | 35.221 | 259.787 | 0.000 | 0.000 | 0.410 | 0.000 | 0.265 | - | 1.530 | WEC this study |
| 2015-08-10 | KT16S313 | 15.823 | 35.171 | 264.727 | 0.000 | 0.000 | 0.308 | 0.000 | 0.260 | - | 2.290 | WEC this study |
| 2015-08-17 | KT16S314 | 16.702 | 35.247 | 267.294 | 0.000 | 0.000 | 0.148 | 0.000 | 0.488 | - | - | WEC this study |
| 2015-08-24 | KT16S315 | 16.584 | 35.162 | 240.779 | 0.000 | 0.000 | 0.300 | 0.050 | 0.570 | - | 0.980 | WEC this study |

| | | | | | | | | | | | | |
|---|---|---|---|---|---|---|---|---|---|---|---|---|
| 2015-08-31 | KT16S316 | 15.847 | 35.101 | 234.150 | 0.833 | 0.140 | 0.617 | 0.133 | 2.028 | - | 1.090 | WEC this study |
| 2015-09-07 | KT16S317 | 15.378 | 35.286 | 234.963 | 1.140 | 0.500 | 0.105 | 0.118 | 1.515 | - | 2.040 | WEC this study |
| 2015-09-14 | KT16S318 | - | - | - | 0.115 | 0.045 | 0.000 | 0.078 | 1.360 | - | 2.510 | WEC this study |
| 2015-09-21 | KT16S319 | - | - | - | 0.450 | 0.150 | 0.145 | 0.110 | 1.850 | - | 1.680 | WEC this study |
| 2015-09-28 | KT16S320 | 15.623 | 35.219 | 236.715 | 0.093 | 0.060 | 0.037 | 0.095 | 2.290 | - | 2.020 | WEC this study |
| 2015-10-05 | KT16S321 | 15.430 | 35.224 | 234.349 | 0.240 | 0.160 | 0.153 | 0.093 | 2.153 | - | 1.010 | WEC this study |
| 2015-10-12 | KT16S322 | 15.200 | 35.228 | 227.175 | 1.225 | 0.735 | 0.308 | 0.165 | 2.510 | - | 0.720 | WEC this study |
| 2015-10-19 | KT16S323 | 14.955 | 35.306 | 231.971 | 1.243 | 0.915 | 0.127 | 0.155 | 1.953 | - | 1.490 | WEC this study |
| 2015-10-26 | KT16S324 | - | - | - | 1.538 | 0.855 | 0.125 | 0.190 | 2.280 | - | 1.180 | WEC this study |
| 2015-11-02 | KT16S325 | 14.617 | 35.302 | 233.253 | 1.990 | 0.530 | 0.123 | 0.233 | 2.193 | - | 0.910 | WEC this study |
| 2015-11-16 | KT16S326 | 13.974 | 35.214 | 238.783 | 4.133 | 0.050 | 0.020 | 0.340 | 3.255 | - | 0.570 | WEC this study |
| 2015-11-23 | KT16S327 | 13.512 | 35.195 | 238.740 | 4.453 | 0.105 | 0.140 | 0.348 | 3.168 | - | 0.520 | WEC this study |
| 2015-12-07 | KT16S328 | 12.597 | 35.255 | 251.008 | 5.758 | 0.050 | 0.350 | 0.453 | 3.257 | - | 0.270 | WEC this study |
| 2015-12-14 | KT16S329 | 12.300 | 35.232 | 252.096 | 6.543 | 0.115 | 0.108 | 0.478 | 3.410 | - | 0.340 | WEC this study |
| 2016-01-04 | KT16S330 | 11.668 | 35.060 | 252.222 | 6.048 | 0.095 | 0.175 | 0.520 | 4.010 | - | 0.320 | WEC this study |
| 2016-01-11 | KT16S331 | 11.124 | 35.064 | 253.197 | 6.098 | 0.075 | 0.063 | 0.485 | 3.888 | - | 0.400 | WEC this study |
| 2016-01-18 | KT16S332 | 10.757 | 35.027 | 254.297 | 7.065 | 0.080 | 0.088 | 0.493 | 3.913 | - | 0.370 | WEC this study |
| 2016-01-25 | KT16S333 | 10.255 | 34.410 | 264.533 | 11.700 | 0.175 | 0.457 | 0.593 | 5.585 | - | 0.500 | WEC this study |
| 2016-02-01 | KT16S334 | - | - | - | 8.250 | 0.070 | 0.130 | 0.528 | 4.425 | - | 0.500 | WEC this study |
| 2016-02-08 | KT16S335 | 10.303 | 35.113 | 259.368 | 7.378 | 0.090 | 0.150 | 0.543 | 3.685 | - | - | WEC this study |
| 2016-02-15 | KT16S336 | 10.204 | 35.107 | 259.207 | 7.095 | 0.090 | 0.000 | 0.515 | 3.455 | - | 0.430 | WEC this study |
| 2016-02-22 | KT16S337 | 9.879 | 34.926 | 265.528 | 8.048 | 0.165 | 0.163 | 0.525 | 3.848 | - | 0.570 | WEC this study |
| 2016-02-29 | KT16S338 | 9.785 | 35.069 | 265.830 | 7.353 | 0.145 | 0.133 | 0.500 | 3.570 | - | 0.540 | WEC this study |
| 2016-03-07 | KT16S339 | 9.451 | 35.013 | 266.768 | 7.473 | 0.235 | 0.135 | 0.473 | 3.663 | - | 0.690 | WEC this study |
| 2016-03-14 | KT16S340 | 9.346 | 34.937 | 277.219 | 6.878 | 0.255 | 0.150 | 0.475 | 3.148 | - | 0.640 | WEC this study |
| 2016-03-21 | KT16S341 | 9.536 | 35.097 | 269.154 | 7.127 | 0.220 | 0.060 | 0.448 | 3.573 | - | 0.950 | WEC this study |
| 2016-03-28 | KT16S342 | 9.460 | 34.970 | 269.982 | 5.705 | 0.200 | 0.355 | 0.430 | 2.910 | - | 1.100 | WEC this study |
| 2016-04-04 | KT16S343 | 9.593 | 35.015 | 270.780 | 4.650 | 0.233 | 0.440 | 0.410 | 1.988 | - | 0.840 | WEC this study |
| 2016-04-11 | KT16S344 | 9.766 | 35.047 | 267.806 | 5.958 | 0.285 | 0.390 | 0.410 | 2.720 | - | 0.610 | WEC this study |
| 2016-04-18 | KT16S345 | 9.999 | 34.928 | 270.794 | 5.170 | 0.180 | 0.165 | 0.333 | 2.580 | - | 1.460 | WEC this study |
| 2016-04-25 | KT16S346 | 10.164 | 34.967 | 276.345 | 5.530 | 0.200 | 0.277 | 0.388 | 2.163 | - | 1.330 | WEC this study |
| 2016-05-02 | KT16S347 | 10.242 | 35.037 | 288.808 | 1.985 | 0.140 | 0.410 | 0.195 | 0.990 | - | 1.360 | WEC this study |
| 2016-05-09 | KT16S348 | 10.635 | 35.166 | 300.819 | 0.180 | 0.025 | 0.190 | 0.103 | 0.655 | - | 2.310 | WEC this study |
| 2016-05-16 | KT16S349 | 11.804 | 35.219 | 305.560 | 0.000 | 0.000 | 0.188 | 0.075 | 0.715 | - | 0.400 | WEC this study |
| 2016-05-23 | KT16S350 | 12.061 | 35.057 | 276.109 | 0.000 | 0.000 | 0.107 | 0.065 | 1.187 | - | 0.300 | WEC this study |
| 2016-05-30 | KT16S351 | 12.783 | 35.165 | 267.811 | 0.010 | 0.005 | 0.263 | 0.045 | 0.935 | - | 0.190 | WEC this study |
| 2016-06-09 | KT16S352 | - | - | - | - | - | - | - | - | - | - | WEC this study |
| 2016-06-20 | KT16S353 | 14.058 | 34.943 | 253.370 | 0.015 | 0.000 | 0.050 | 0.028 | 0.238 | - | 0.520 | WEC this study |
| 2016-06-27 | KT16S354 | 13.810 | 35.166 | 256.653 | 0.013 | 0.000 | 0.087 | 0.015 | 0.563 | - | 0.870 | WEC this study |
| 2016-07-04 | KT16S355 | 14.458 | 35.016 | 258.718 | 0.020 | 0.000 | 0.127 | 0.000 | 0.538 | - | 0.880 | WEC this study |
| 2016-07-11 | KT16S356 | 14.652 | 35.158 | 254.262 | 0.000 | 0.000 | 0.090 | 0.015 | 1.260 | - | 1.130 | WEC this study |
| 2016-07-18 | KT16S357 | 16.244 | 35.222 | 253.382 | 0.000 | 0.000 | 0.190 | 0.025 | 1.200 | - | 0.480 | WEC this study |
| 2016-07-25 | KT16S358 | 16.760 | 35.253 | 279.705 | 0.000 | 0.000 | 0.195 | 0.000 | 1.043 | - | 1.530 | WEC this study |
| 2016-08-01 | KT16S359 | 16.125 | 35.235 | 255.200 | 0.020 | 0.000 | 0.093 | 0.050 | 1.108 | - | 1.280 | WEC this study |
| 2016-08-08 | KT16S360 | 15.544 | 35.139 | 260.108 | 0.000 | 0.000 | 0.018 | 0.010 | 1.655 | - | 0.860 | WEC this study |

| | | | | | | | | | | | | |
|---|---|---|---|---|---|---|---|---|---|---|---|---|
| 2016-08-15 | KT16S361 | 16.232 | 35.213 | 265.939 | 0.020 | 0.000 | 0.020 | 0.000 | 1.550 | - | 0.840 | WEC this study |
| 2016-08-22 | KT16S362 | 16.651 | 35.197 | 245.543 | 0.005 | 0.000 | 0.218 | 0.040 | 1.890 | - | 3.840 | WEC this study |
| 2016-08-29 | KT16S363 | 16.891 | 35.273 | 234.857 | 0.030 | 0.015 | 0.225 | - | 1.207 | - | 1.050 | WEC this study |
| 2016-09-05 | KT16S364 | 16.483 | 35.178 | 228.534 | 0.303 | 0.160 | 0.457 | 0.095 | 1.438 | - | 1.510 | WEC this study |
| 2016-09-12 | KT16S365 | 17.079 | 35.116 | 234.464 | 0.030 | 0.020 | 0.400 | 0.040 | 0.883 | - | 1.400 | WEC this study |
| 2016-09-19 | KT16S366 | 16.253 | 35.101 | 177.332 | 0.655 | 0.190 | 0.133 | 0.103 | 1.373 | - | 2.500 | WEC this study |
| 2016-09-26 | KT16S367 | 16.272 | 35.152 | 229.280 | 0.790 | 0.265 | 0.318 | 0.145 | 1.465 | - | 1.530 | WEC this study |
| 2016-10-10 | KT16S368 | 15.803 | 35.283 | 228.833 | 1.345 | 0.520 | 0.108 | 0.150 | 1.388 | - | 1.590 | WEC this study |
| 2016-10-17 | KT16S369 | 15.285 | 35.281 | 225.997 | 1.565 | 0.665 | 0.388 | 0.185 | 1.648 | - | - | WEC this study |
| 2016-10-24 | KT16S370 | - | - | - | 1.095 | 0.820 | 0.353 | 0.210 | 1.455 | - | 0.940 | WEC this study |
| 2016-10-31 | KT16S371 | 15.240 | 35.272 | 228.443 | 1.488 | 1.060 | 0.258 | 0.235 | 1.720 | - | 0.610 | WEC this study |
| 2016-11-07 | KT16S372 | 14.771 | 35.306 | 229.158 | 1.995 | 1.160 | 0.148 | 0.250 | 1.820 | - | 0.950 | WEC this study |
| 2016-11-14 | KT16S373 | 14.136 | 35.277 | 229.580 | 2.650 | 1.317 | 0.140 | 0.268 | 2.243 | - | 0.800 | WEC this study |
| 2016-11-21 | KT16S374 | 12.517 | 34.469 | 239.213 | - | - | - | - | - | - | 1.090 | WEC this study |
| 2016-11-28 | KT16S375 | 12.877 | 35.172 | 233.036 | 3.905 | 0.540 | 0.127 | 0.323 | 2.695 | - | 0.680 | WEC this study |
| 2016-12-05 | KT16S376 | 12.128 | 35.077 | 240.751 | 4.640 | 1.385 | 0.018 | 0.488 | 4.673 | - | 0.630 | WEC this study |
| 2016-12-12 | KT16S377 | 12.276 | 35.155 | 241.253 | 4.175 | 0.075 | 0.100 | 0.350 | 3.630 | - | 0.510 | WEC this study |
| 2016-12-19 | KT16S378 | 12.300 | 35.097 | 199.094 | 4.078 | 0.075 | 0.123 | 0.380 | 3.440 | - | 0.630 | WEC this study |
| 2017-01-02 | KT16S379 | 11.628 | 35.237 | 244.892 | - | - | - | - | - | - | 0.550 | WEC this study |
| 2017-01-09 | KT16S380 | 10.924 | 35.089 | 247.747 | 6.103 | 0.228 | 0.155 | 0.428 | 3.633 | - | 0.510 | WEC this study |
| 2017-01-16 | KT16S381 | 10.670 | 35.037 | 247.367 | 5.983 | 0.270 | 0.100 | 0.428 | 3.838 | - | 0.620 | WEC this study |
| 2017-01-23 | KT16S382 | 10.380 | 35.171 | 251.196 | 5.543 | 0.178 | 0.000 | 0.413 | - | - | 0.800 | WEC this study |
| 2017-01-30 | KT16S383 | 10.165 | 35.202 | 254.132 | 5.433 | 0.190 | 0.083 | 0.420 | 3.388 | - | 0.670 | WEC this study |
| 2017-02-06 | KT16S384 | 9.622 | 34.744 | 256.172 | 7.735 | 0.323 | 0.313 | 0.490 | 4.660 | - | 0.670 | WEC this study |
| 2017-02-13 | KT16S385 | - | - | - | 5.867 | 0.180 | 0.095 | 0.455 | 3.305 | - | 1.320 | WEC this study |
| 2017-02-20 | KT16S386 | 9.756 | 34.861 | 261.753 | 8.048 | 0.265 | 0.178 | 0.448 | 3.708 | - | 0.810 | WEC this study |
| 2017-02-27 | KT16S387 | 9.706 | 35.199 | 255.756 | 5.658 | 0.120 | 0.000 | 0.455 | 3.140 | - | 1.190 | WEC this study |
| 2017-03-06 | KT16S388 | 9.675 | 35.174 | 256.202 | 5.653 | 0.165 | - | 0.425 | 2.750 | - | 0.800 | WEC this study |
| 2017-03-13 | KT16S389 | 9.942 | 34.780 | 261.974 | 7.593 | 0.258 | - | 0.435 | 3.613 | - | 0.460 | WEC this study |
| 2017-03-20 | KT16S390 | 10.158 | 35.149 | 217.058 | 5.695 | 0.280 | 0.163 | 0.410 | 2.650 | - | 0.490 | WEC this study |
| 2017-03-27 | KT16S391 | - | - | - | 5.783 | 0.340 | 0.145 | 0.408 | 2.863 | - | 0.860 | WEC this study |
| 2017-04-03 | KT16S392 | 10.717 | 35.125 | 266.164 | 5.123 | 0.285 | 0.125 | 0.355 | 2.543 | - | 1.930 | WEC this study |
| 2017-04-10 | KT16S393 | 10.922 | 35.015 | 277.635 | 3.888 | 0.190 | 0.093 | 0.238 | 2.460 | - | 2.990 | WEC this study |
| 2017-04-17 | KT16S394 | - | - | - | 2.427 | 0.190 | 0.350 | 0.253 | 1.763 | - | 1.330 | WEC this study |
| 2017-04-24 | KT16S395 | 11.702 | 35.163 | 290.605 | 0.608 | 0.055 | 0.355 | 0.145 | 0.988 | - | 2.100 | WEC this study |
| 2017-05-01 | KT16S396 | 10.999 | 35.230 | 264.882 | 0.883 | 0.090 | 0.810 | 0.225 | 0.520 | - | 1.000 | WEC this study |
| 2017-05-08 | KT16S397 | 11.514 | 35.277 | 278.736 | 0.187 | 0.040 | 0.170 | - | 0.273 | - | 0.800 | WEC this study |
| 2017-05-15 | KT16S398 | 12.336 | 34.981 | 259.053 | 0.000 | 0.000 | 0.000 | 0.103 | 0.233 | - | 0.760 | WEC this study |
| 2017-05-22 | KT16S399 | 12.971 | 35.050 | 261.086 | 0.000 | 0.000 | 0.000 | 0.050 | 0.158 | - | 1.750 | WEC this study |
| 2017-05-29 | KT16S400 | 13.730 | 35.131 | 266.637 | 0.000 | 0.000 | 0.020 | 0.015 | 0.097 | - | 2.680 | WEC this study |
| 2017-06-12 | KT16S401 | 13.438 | 35.060 | 250.878 | 0.028 | 0.000 | 0.183 | 0.070 | 0.400 | - | 0.650 | WEC this study |
| 2017-06-19 | KT16S402 | 16.290 | 35.207 | 253.371 | 0.000 | 0.000 | 0.148 | 0.000 | 0.133 | - | 0.890 | WEC this study |
| 2017-06-26 | KT16S403 | 15.288 | 35.120 | 197.268 | 0.000 | 0.000 | 0.133 | 0.030 | 0.340 | - | 0.910 | WEC this study |
| 2017-07-03 | KT16S404 | 14.962 | 35.263 | 252.908 | 0.000 | 0.000 | - | 0.035 | 0.145 | - | 0.390 | WEC this study |
| 2017-07-17 | KT16S405 | 16.141 | 35.070 | 210.064 | 0.040 | 0.000 | - | 0.040 | 0.433 | - | 0.380 | WEC this study |

| | | | | | | | | | | | | |
|---|---|---|---|---|---|---|---|---|---|---|---|---|
| 2017-07-24 | KT16S406 | 16.642 | 35.110 | 231.812 | 0.190 | 0.035 | - | 0.068 | 1.127 | - | 1.080 | WEC this study |
| 2017-07-31 | KT16S407 | - | - | - | 0.015 | 0.005 | - | 0.048 | 1.250 | - | - | WEC this study |
| 2017-08-07 | KT16S408 | 16.100 | 35.173 | 174.591 | 0.000 | 0.000 | - | 0.000 | 0.433 | - | - | WEC this study |
| 2017-08-14 | KT16S409 | 16.174 | 35.256 | 231.278 | 0.000 | 0.000 | - | 0.015 | 0.650 | - | 0.890 | WEC this study |
| 2017-08-21 | KT16S410 | 17.698 | 35.295 | 233.601 | 0.015 | 0.000 | - | 0.000 | 0.818 | - | 0.970 | WEC this study |
| 2017-08-28 | KT16S411 | 16.400 | 35.143 | 222.646 | 0.000 | 0.000 | - | 0.000 | 0.885 | - | 0.390 | WEC this study |
| 2017-09-04 | KT16S412 | 15.213 | 34.678 | 220.830 | 0.000 | 0.000 | - | 0.035 | 1.168 | - | 0.900 | WEC this study |
| 2017-09-18 | KT16S413 | 15.491 | 35.051 | 178.634 | 0.993 | 0.730 | 0.130 | 0.155 | 1.795 | - | 1.360 | WEC this study |
| 2017-09-25 | KT16S414 | 15.567 | 35.028 | 225.872 | 1.950 | 0.750 | 0.105 | 0.185 | 2.700 | - | 1.140 | WEC this study |
| 2017-10-30 | KT16S415 | 14.605 | 35.191 | 224.027 | 3.295 | 0.260 | 0.060 | 0.333 | 3.410 | - | 1.220 | WEC this study |
| 2017-11-13 | KT16S416 | 14.048 | 35.256 | 225.475 | 3.575 | 0.105 | 0.030 | 0.348 | 3.195 | - | 0.920 | WEC this study |
| 2017-11-20 | KT16S417 | 13.515 | 35.121 | 224.565 | 4.225 | 0.125 | 0.083 | 0.360 | 3.670 | - | 0.960 | WEC this study |
| 2017-12-04 | KT16S418 | 12.526 | 35.199 | 232.053 | 4.447 | 0.098 | 0.130 | 0.380 | 3.770 | - | 0.830 | WEC this study |
| 2017-12-11 | KT16S419 | 11.226 | 34.901 | 184.210 | 7.395 | 0.225 | 0.320 | 0.440 | 5.020 | - | 0.800 | WEC this study |
| 2017-12-18 | KT16S420 | 10.752 | 34.792 | 241.412 | 7.345 | 0.225 | 0.127 | 0.438 | 4.775 | - | 0.710 | WEC this study |
| 2018-01-08 | KT16S421 | 10.307 | 35.048 | - | 7.040 | 0.110 | 0.147 | 0.490 | 3.893 | - | 0.530 | WEC this study |
| 2018-01-22 | KT16S422 | 9.970 | 34.681 | - | 8.433 | 0.160 | 0.228 | 0.503 | 4.010 | - | 0.440 | WEC this study |
| 2018-01-29 | KT16S423 | 9.982 | 35.000 | - | 7.540 | 0.110 | 0.138 | 0.475 | 3.408 | - | 0.580 | WEC this study |
| 2018-02-05 | KT16S424 | 9.608 | 34.974 | 236.476 | 7.260 | 0.135 | 0.170 | 0.483 | 3.418 | - | 0.510 | WEC this study |
| 2018-02-12 | KT16S425 | 8.873 | 34.885 | - | 7.563 | 0.185 | 0.160 | 0.450 | 3.063 | - | 0.550 | WEC this study |
| 2018-02-19 | KT16S426 | 9.125 | 34.270 | - | 11.287 | 0.270 | 0.453 | 0.453 | 4.653 | - | 1.290 | WEC this study |
| 2018-02-26 | KT16S427 | 8.487 | 35.100 | - | 10.110 | - | - | - | 2.000 | - | 1.080 | WEC this study |
| 2018-03-05 | KT16S428 | 7.787 | 35.046 | 232.564 | 6.938 | 0.248 | 0.207 | 0.453 | 3.148 | - | 0.420 | WEC this study |
| 2018-03-12 | KT16S429 | - | - | - | 12.980 | 0.300 | 1.035 | 0.523 | 5.718 | - | 0.350 | WEC this study |
| 2018-03-19 | KT16S430 | 7.849 | 34.647 | 243.424 | 8.678 | 0.235 | 0.385 | 0.475 | 3.715 | - | 0.950 | WEC this study |
| 2018-04-02 | KT16S431 | 8.007 | 34.594 | 248.062 | 8.745 | 0.198 | 0.353 | 0.435 | 3.810 | - | 1.020 | WEC this study |
| 2018-04-09 | KT16S432 | 8.366 | 34.622 | 244.833 | 15.668 | 0.275 | 0.360 | 0.355 | 6.413 | - | 3.950 | WEC this study |
| 2018-04-19 | KT16S433 | - | - | - | - | - | - | - | - | - | - | WEC this study |
| 2018-04-26 | KT16S434 | - | - | - | - | - | - | - | - | - | - | WEC this study |
| 2018-04-30 | KT16S435 | - | - | - | 0.840 | 0.060 | 0.985 | 0.100 | 0.493 | - | - | WEC this study |
| 2018-05-07 | KT16S436 | 11.559 | 35.064 | - | 0.025 | 0.000 | 0.080 | 0.035 | 0.338 | - | 1.450 | WEC this study |
| 2018-05-14 | KT16S437 | 11.513 | 34.920 | - | 0.000 | 0.000 | 0.125 | 0.000 | 0.233 | - | 0.190 | WEC this study |
| 2018-05-21 | KT16S438 | 11.818 | 35.188 | - | 0.000 | 0.000 | 0.053 | 0.068 | 0.475 | - | 0.380 | WEC this study |
| 2018-05-30 | KT16S439 | - | - | - | - | - | - | - | - | - | - | WEC this study |
| 2018-06-04 | KT16S440 | 14.526 | 35.150 | - | 0.000 | 0.000 | 0.000 | 0.015 | 0.330 | - | 0.250 | WEC this study |
| 2018-06-18 | KT16S441 | 13.838 | 35.073 | - | 0.037 | 0.000 | 0.143 | 0.050 | 0.160 | - | 0.730 | WEC this study |
| 2018-06-25 | KT16S442 | 15.283 | 35.150 | - | 0.020 | 0.000 | 0.115 | 0.040 | 0.073 | - | 0.350 | WEC this study |
| 2018-07-02 | KT16S443 | 18.153 | 35.112 | 208.245 | 0.040 | 0.000 | 0.250 | 0.013 | 0.085 | - | 0.620 | WEC this study |
| 2018-07-09 | KT16S444 | 18.148 | 35.115 | 207.566 | 0.033 | 0.000 | 0.128 | 0.033 | 0.063 | - | 0.290 | WEC this study |
| 2018-07-16 | KT16S445 | 19.167 | 35.017 | 177.404 | 0.000 | 0.000 | 0.000 | 0.000 | - | - | 0.230 | WEC this study |
| 2018-07-23 | KT16S446 | 16.853 | 35.097 | 201.752 | 0.000 | 0.000 | 0.000 | 0.030 | - | - | 0.590 | WEC this study |
| 2018-07-30 | KT16S447 | 16.724 | 35.086 | 245.903 | 0.000 | 0.000 | 0.185 | 0.093 | - | - | 1.090 | WEC this study |
| 2018-08-06 | KT16S448 | 17.884 | 34.964 | 265.454 | 0.020 | 0.000 | 0.250 | 0.020 | 0.568 | - | 0.380 | WEC this study |
| 2018-08-13 | KT16S449 | 16.459 | 34.967 | 267.668 | 0.000 | 0.000 | 0.000 | 0.050 | 0.950 | - | 2.150 | WEC this study |
| 2018-08-20 | KT16S450 | 16.471 | 34.966 | 266.252 | 0.015 | 0.000 | 0.440 | 0.040 | 1.613 | - | 1.050 | WEC this study |

| | | | | | | | | | | | | |
|---|---|---|---|---|---|---|---|---|---|---|---|---|
| 2018-08-27 | KT16S451 | 16.208 | 34.962 | 216.404 | 0.010 | 0.000 | 0.255 | 0.023 | 1.580 | - | 2.320 | WEC this study |
| 2018-09-03 | KT16S452 | 16.675 | 35.040 | 244.267 | 0.050 | 0.020 | 0.755 | 0.135 | 1.700 | - | 0.640 | WEC this study |
| 2018-09-17 | KT16S453 | - | - | - | 0.000 | 0.000 | - | 0.053 | 2.030 | - | - | WEC this study |
| 2018-12-03 | KT16S454 | 12.117 | 34.628 | 253.638 | 7.393 | 0.243 | 0.305 | 0.445 | 4.398 | - | 0.760 | WEC this study |
| 2018-12-10 | KT16S455 | 11.572 | 33.884 | 258.299 | 11.873 | 0.227 | 0.440 | 0.573 | 5.833 | - | 0.570 | WEC this study |
| 2018-12-17 | KT16S456 | - | - | - | 8.525 | 0.188 | 0.388 | 0.493 | 4.558 | - | - | WEC this study |
